# Aquaporin-4 mediated aggregation of Alzheimer’s amyloid β-peptide

**DOI:** 10.1101/2023.02.08.527707

**Authors:** Nikhil Maroli

**Affiliations:** Computational Biology Division, DRDO Center for Life Science, Bharathiar University Campus, Coimbatore 641046, Tamil Nadu, India

**Keywords:** Alzheimers disease, Amyloid oligomers, Aquaporin-4, amyloid aggregation, amyloid clearance, amyloid peptide 42

## Abstract

Clearance of Alzheimer’s amyloid oligomers from the brain is crucial for preventing cell toxicity. Dementia complications arise as a result of apoptosis, which is caused by peptide plaques on the lipid surface of cells. Here, we employed all-atom and coarse-grained molecular dynamics simulations to investigate the aggregation of amyloid peptides at the lipid surface and the role of the Aquaporin-4 (AQP4) in facilitating peptide clearance from astrocytes. The network of protein-protein interactions through text mining revealed that the expression of AQP4 and amyloid aggregation were strongly correlated. It has also been revealed that the role of aquaporins in the etiology of Alzheimer’s involves several interconnected proteins and pathways. The nature of aggregation at the surface of the 1-palmitoyl-2-oleoyl-sn-glycero-3-phosphocholine (POPC) lipid bilayer was revealed by the interaction of amyloid oligomers. The membrane-bound pore region of AQP4 interacts with the peptide and slows its aggregation. This interaction maintains the helical content of the peptide while lowering its toxicity at the lipid surface. The hydrophobicity of the peptide also decreased because of these interactions, which may help in the removal of the peptide from astrocytes. Long-term coarse-grained MD simulations demonstrated different features of oligomer aggregation at the surface and strong oligomer attraction to AQP4, which inhibited aggregation. Additionally, the water dynamics of aquaporins demonstrate how the selectivity filter is broken to disrupt water flow. Our findings also provide insight into the physiological alterations in brain tissue associated with Alzheimer’s disease, including water retention and increased water flow in the CSF. Furthermore, in vitro thioflavin fluorescence spectroscopy revealed a slower aggregation of the peptide in the presence of AQP4.

## Introduction

The blood-brain barrier (BBB) plays an important role in the transportation and neuronal signaling of molecules. The BBB, formed by endothelial cells that line cerebral microvessels, is a highly permeable ’barrier’ between the circulating blood and the central nervous system. The BBB is composed of cerebral blood vessels, pericytes, perivascular microglia, perivascular neurons and astrocytes [1–2]. Astrocytes are star-shaped glial cells that impinge on the basement membrane of blood vessels and are highly enriched in water and potassium channels. One potential role of astrocytes is to balance potassium transport and water movement via potassium and aquaporin-4, situated in the endfeet [3–7]. Transporting therapeutic candidates across the BBB is a challenging and crucial aspect of brain-related disease. Among these, Alzheimer’s disease, a common cause of dementia, has attracted the attention of researchers in the past few decades [8]. A neurodegenerative disease starts with short-term memory loss, language problems, and gradual loss of bodily functions leading to death [9–10]. The disease is progressive, and abnormalities are first detected in the frontal and temporal lobes of the brain, and then slowly progress to other areas of the brain depending on the individual. It is associated with the deposition of amyloid-β plaques in the extracellular spaces and walls of blood vessels [11–12]. In addition, aggregation of the microtubule protein tau in neurofibrillary tangles causes memory loss and confusion, leading to loss of cognitive power. Amyloid-β(Aβ) peptides are the main component of senile plaques, and several pieces of evidence support that Aβ is a pathogenic peptide in Alzheimer’s disease, which is derived from the cleavage of a larger glycoprotein-amyloid precursor protein (APP) [13–14]. APP plays a significant role in a range of biological functions such as neuronal development, signalling, and intracellular transport [15–16]. APP plays a major role in Alzheimer’s disease pathogenesis and is expressed in many tissues and neuronal synapses. This precursor protein is cut by β-secretases and γ-secretases to 37–49 amino acid residue peptides called Aβ [17–18]. APP is cleaved through a non-amyloidogenic or amyloidogenic pathway, generating α-or β-C-terminal fragments. Later, this is further cleaved by γ-secretase to liberate the P3 and Aβ peptides. The C-terminal fragment of APP produced by β-secretase is further cleaved by γ-secretase at multiple sites to produce the main final form of Aβ which contains 37-49 amino acids [18]. Aβ monomers aggregate into various forms such as oligomers, protofibrils, and amyloid fibrils. Amyloid fibrils are insoluble and form plaques in the brain, whereas oligomers are soluble and are distributed throughout the brain [19–20]. Glenner et al. first discovered the amyloid-β from an extracellular deposit, though it’s an intrinsically unstructured peptide and difficult to crystalize [21]. Later, the three-dimensional solution structure of different fragments of Aβ was determined by NMR and molecular dynamics techniques; NMR also suggested different conformational states for Aβ_1-40_ and Aβ_1-42_ [22]. The residues 31-34 and 38-41 form a β-hairpin that reduces the C-terminal flexibility of Aβ_42_, which is responsible for the higher tendency of Aβ_42_ than Aβ_40_ to form amyloid plaques [23]. Yang et al. used the replica exchange molecular dynamics method to suggest Aβ_40_ and Aβ_42_ can indeed populate multiple discrete conformations, including α-helix or β-sheet and they observed rapid transition in these conformations [24]. Sgourakis et al. used statistical analysis along with molecular dynamics simulations to identify a multiplicity of discrete conformational clusters in Aβ_42_ [25]. Amyloid-β monomers can form higher-order assemblies from low-molecular-weight oligomers, such as dimers, trimers, tetramers, pentamers, and high-molecular-weight oligomers, such as nonamers and dodecamers, to protofibrils and fibrils. Given the difficulty of obtaining high-resolution structures, alternative theoretical and simulation approaches have been used to rationalize the physiochemical properties of amyloid-β and plaque formation.

Alzheimer’s diseases are an imbalance between Aβ production and clearance, for an efficient strategy to cure AD it is essential to understand the clearance systems in the brain, especially transport across the blood-brain barrier. It is suggested that the astroglial-mediated interstitial fluid (ISF) bulk flow, known as the glymphatic system might be contributed in a larger portion towards the clearance of water-soluble Aβ [26–32]. Donna et al. showed that vascular amyloid deposition in AD and cerebral amyloid angiopathy (CAA) in both mouse and human models results in the mislocalization of aquaporin-4 expression and changes in the expression of specific potassium channels, both of which are essential for maintaining brain physiology [33–34]. The animal lacks aquaporin-4 in astrocytes and exhibits slow cerebrospinal fluid (CSF) influx through the brain parenchyma and perivascular spaces. Fluorescent-tagged amyloid-β is transported along this route, and deletion of AQP4 genes suppresses soluble amyloid-β transportation. The expression of AQP4 in the brains of patients with or without CAA showed impaired water transport. Previous studies have suggested that the expression of aquaporin-4 is high in the brain affected by edema and cerebral ischemia [35]. Furthermore, recent studies have shown that AQP4 is a critical target in neuromyelitis optica [36]. The amyloid-associated proteins α1-antichuumotrypsim, apo-protein, and C1q modify amyloid-β uptake by astrocytes. The presence of intracellular Aβ in human astrocytes has been reported by several researchers [29]. However, the mechanism of Aβ uptake and degradation in these cell lines has not yet been studied in detail. Nielsen et al. suggested that the astrocytes have the potential to bind and ingest Aβ [37]. They also showed that scavenger receptor class B type 1 (SCARB1) plays an important role in Aβ binding and endocytosis in adult mouse astrocytes. Furthermore, the expression of IDE (Insulin-degrading enzyme), NEP (Neutral endopeptidase), SCARB1 (Scavenger receptor class B type 1), MARCO (Macrophage receptor with collagenous structure), LRP2 (Low-density lipoprotein receptor-related protein 2), and MMP-9 (Matrix metalloproteinase-9) associated with Aβ clearance, was studied by Sandra et al. [26]. They proposed that dysfunctional regulation of gene expression in AD astrocytes leads to defective clearance of Aβ and promotes the accumulation of Aβ in the brain.

Several computational studies have dealt with amyloid-β aggregation and the inhibition of the aggregation process via various small molecules; however, the underlying inhibitory mechanism remains elusive. Kolandaivel et al. studied the role of various metal ions and the heme complex in the brain in action with the Aβ monomer [38]. Several reports have shown the inhibition of amyloid-β aggregation by analyzing secondary structural changes in the Aβ peptide. Only a few researchers have focused on the interactions of Aβ oligomers and plaques with their surrounding environment. Davis et al. studied the interaction of amyloid-β with phospholipid bilayers and tetramer formation using molecular dynamic simulations [39]. They studied the Aβ_1-42_ peptide interaction with zwitterion DPPC lipids and anionic DOPS phospholipid bilayers and found that the Aβ was attracted towards these membrane layers and increased the peptide-peptide interaction, leading to fast aggregation. They provide further information on the aggregation pathway of Aβ and the putative toxic mechanism in the pathogenesis of Alzheimer’s disease. Urbanc et al. studied amyloid-β dimer formation using discrete and coarse-grained molecular dynamics (CGMD) simulations to estimate the thermodynamic stability of all dimer conformations [40]. The dimer conformations possess higher free energy differences than the monomer, and they compare the free energy between the Aβ_1-42_ and Aβ_1-40_ dimers. Furthermore, they found that Aβ oligomerization is not accompanied by the formation of thermodynamically stable planar β-strand dimers. The aggregation of peptides on the lipid bilayer using CGMD simulation in an implicit solvent showed that amyloid-β self-assembles on the surface into β-rich fibrillar aggregates, even if they are dissolved in a disordered oligomer form [39]. The key interaction between the oligomerization is the Asn-Asn interaction which works like “glue” by sticking the DFNKF strands together. A CGMD study of fibril-forming amphipathic peptides in the presence of lipid vesicles showed that the deposition of protein aggregates on the cell membrane results in leakage and cell disruption.

Aquaporin-4 is a hetero-transmembrane protein that is mostly expressed in astrocyte endfeet. The blood-brain barrier maintains important brain activities such as the release or uptake of neurotransmitters. These are made possible by astrocytes and the distinctive star-shaped glial cells found in the brain and spinal cord. We hypothesized that the interaction between amyloid-β and aquaporin-4 is crucial for the removal of amyloid-β from the brain. Furthermore, the altered brain physiology observed in previous studies suggests a possible strong interaction between these proteins and peptides. To gain insight into the atomic-level interactions between amyloid-β and aquaporin-4, we conducted an all-atom molecular dynamics simulation of amyloid-β on the surfaces of lipid bilayers (POPC) and AQP4 embedded in POPC. Coarse-grained molecular dynamics simulations were used to examine the interaction of oligomers and the aggregation process at the lipid and lipid-protein surfaces. Additionally, thioflavin fluorescence spectroscopy was used to assess aggregation. Finally, protein-protein network analysis through text mining and aquaporin-4 water dynamics demonstrated a significant association between amyloid peptide aggregation.

## Result and Discussions

The structural changes in amyloidogenic peptides in the presence of aquaporin-4 and the lipid bilayer were analyzed using a 200 ns all-atom molecular dynamics simulation. Structural features such as secondary structure, salt bridge distances, hydrogen bonds, and interaction energies were exclusively calculated. The salt bridge and hydrophobicity of peptides have a significant influence on conformational changes, aggregation, and secondary structural changes. On the bilayer surface, lipid characteristics and conformational changes in the peptide were simulated using coarse-grained molecular dynamics simulations. In addition, thioflavin fluorescence spectroscopy was used to assess peptide aggregation in *vitro*. Furthermore, the protein-protein interaction network through text mining revealed hidden pathways involved in AQP4-Aβ clearance in the brain. The water dynamics of AQP4 provide insight into the brain tissue changes induced by Alzheimer’s disease.

### Characterization of peptide conformation at the surface of the lipid bilayer and AQP4 - all atom MD simulation

The 42 amino acids peptide contains three important regions (Figure 1a) that are crucial for the aggregation process: the central hydrophobic core region (Leu17-Ala21), loop region (Asp23-Lys28), and second hydrophobic region (Gly29-Met35) [41–44]. The native amyloid peptide (Figure 1a) monomer secondary structure consisted of 79.4 ± 0.8 % helices and 20.6 ± 0.8 % coils. The residues Asp1-Ser8 adopt the bend region and Gly9-Ser26 show a long α-helix. The interaction between His6-Gly9, Asp7-Gly9, Ala2-Arg5, and Tyr10-Gln15 residues helps stabilize the β-hairpin structure in the region Asp1-Gln15. To elucidate secondary structural transitions, a single peptide was simulated on the surface of the POPC lipid bilayer. During simulation, the monomer unit of amyloid-β exhibited a considerable reduction in its helical conformation; the change was gradual, and the final frame comprised 36.8 ± 1.1 % helices and 62.1 ± 1.1% coils (Figure 2a). Figure 2(a-f) depicts the time-dependent changes in the secondary structure of the peptide in both lipids and AQP4. To prevent initial interactions between the peptide and lipid molecules, we positioned the peptide 2.5 nm away from the surface of the lipid bilayer in an aqueous environment. The transition from α-helix to a β-sheet-rich structure is a critical part of amyloidogenic oligomerization. We have embedded aquaporin-4 in the lipid membrane and kept the amyloid-β at a distance of 2.5 nm from its surface. The Dictionary of Protein Secondary Structure (DSSP) calculation of the monomer secondary structure at the AQP4 surface revealed a transition (first frame to last frame) from 79.4 ± 0.8 % helices and 20.6 ± 0.8 % to 77.2 ± 1% helix and 23.8± 1 % coil. The helical content of the peptide was retained in the presence of aquaporin-4 in the membrane. At the lipid surface, Asp1-Phe19, Lys28-Ile31, and Val39-Ala42 exhibited a coiled conformation, whereas Phe20-Asn27 and Ile32-Gly38 showed helical structures. In contrast, the interaction of AQP4 revealed a helix at Ala2-Gly25 and Ile3-Gly38, and a coil at Ser26-Ala30. Furthermore, the average distance between the COO^-^ and NH_3_^+^ moieties of the residues was used to compute the salt-bridge distance between Asp23-Lys28 and Glu22-Lys28 (Figure 3a-b). The Asp23-Lys28 salt bridge plays a crucial role in fibril formation. The direct salt bridge appeared at 4.3 Å, whereas the indirect or water-mediated salt bridge appeared between 4.3 Å and 7.0 Å [38]. Additionally, indirect salt bridges of 12 Å and 11 Å were observed, indicating weak intramolecular interactions between these residues [38]. Additionally, these interactions in the region Glu22-Lys28 preserve the turn structure throughout the simulation. When Asp23 and Lys28 are separated by a significant distance, the peptide forms a β-hairpin with a turn between residues Ala21-Val24. However, the interaction of the lipids revealed a coiled conformation. In addition, the stability of the Asp23-Lys28 salt bridge was assessed by measuring the electrostatic energies between the negatively charged Asp23 residues and positively charged Lys28 residues (Figure 3c). In the first 50 ns, an unfavorable electrostatic interaction between Asp23 and Lys28 was observed. This is due to the interaction of the POPC lipid headgroups that destabilize the peptide salt bridge. At the POPC surface, the peptide is exposed to a predominantly hydrophobic environment, which leads to secondary structural changes. However, the interaction of AQP4 with the peptide in the first 80 ns showed less unfavorable interaction, which arises from the lipid and charged group interaction from AQP4. In addition, we assessed the salt bridge distance and electrostatic interaction energy between Glu22-Lys28 residues (Figure 3d). Up to 40 ns, there were no interactions between these residues in the peptide, and it was later found to show favorable electrostatic interactions that stabilize the salt bridge. It is also noted from Figure 2a, that these residues stabilize the loop region, and salt bridges can affect the peptide’s conformational stability, aggregation propensity, and toxicity. Additionally, Figure 4a-c depicts the number of hydrogen bonds formed between the peptides and the lipid bilayer, which were estimated assuming an acceptor-hydrogen distance of less than 3 Å and a donor-hydrogen-acceptor angle greater than 120 °. There were an average of four hydrogen bonds between AQP4 and the monomer peptide, and none between the lipid headgroups and monomer peptide. Furthermore, Figure 4d-i illustrates the non-bonded interaction energies such as electrostatic and van der Waals between the monomers and lipids in the presence and absence of AQP4. The interaction of monomers with lipids is crucial because it determines lipid toxicity and conformational changes of the peptide. As depicted in Figure 4d-i, a significant amount of electrostatic interaction energy existed between the monomer and the lipids. However, the presence of aquaporin in the membrane reduced the non-bonded interaction energy between the monomer and lipids. Consequently, these peptides exhibit strong electrostatic and van der Waals interactions with AQP4. The average electrostatic interaction energy between the peptides and membrane is found to be −336.5±13.6 kJ/mol, whereas the average van der Waals interaction energy is −239.45±4.8 kJ/mol. However, the presence of aquaporin in the membrane reduced and found to be an average of -46.09±8.6 kJ/mol and -14.6±2.2 kJ/mol electrostatic and van der Waals interactions energies respectively. In addition, the distribution of the peptide monomer on the POPC lipid bilayers was more extensive than that of AQP4. The electrostatic interaction between the monomer and lipid found to be increase after 120 ns due to the movement of the peptide at the surface (Figure 4f). This also enhances the peptide induced lipid toxicity.

**Figure 1.**
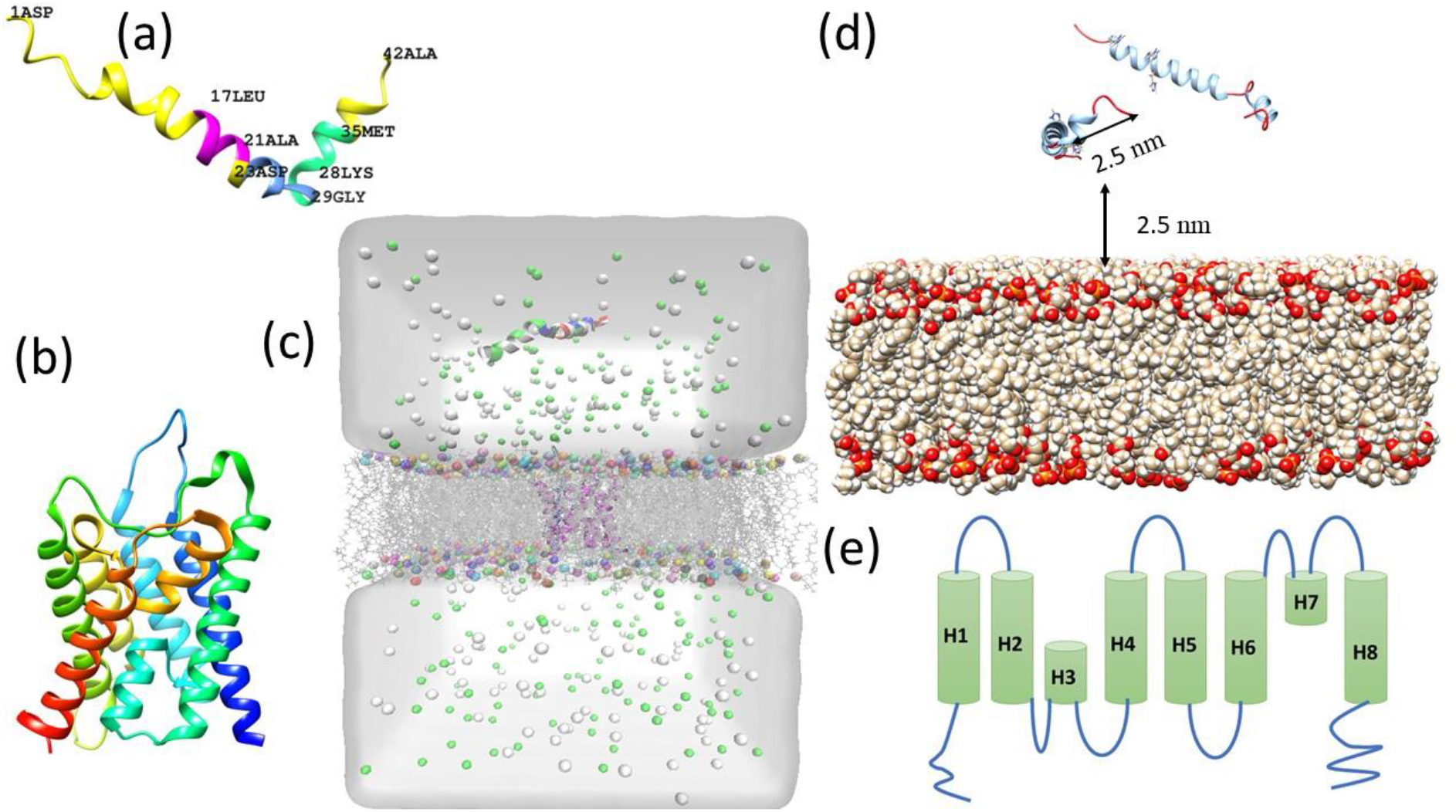
The simulation details are as follows: (a) represents the NMR structure of the amyloid-β peptide. The central hydrophobic core region (Leu17-Ala21), loop region (Asp23-Lys28) and second hydrophobic region (Gly29-Met35) are presented in different colors. (b) X-ray crystal structure of the aquaporin-4 monomer unit and (c) simulation system comprising a POPC lipid bilayer, water boxes, ions, and aquaporin monomer units. Water is represented as a continuous white box, ions are represented in green and white balls, aquaporin is represented as ribbon, and lipids are represented as light grey sticks. (d) represents the initial position of amyloidogenic dimer at the surface of the POPC lipid bilayer. Monomers were separated by a distance of 2.5 nm. (e) Representation of the aquaporin-monomer secondary structures. The helices were represented as H1-H8 and monomer unit is provided for better clarity.

**Figure 2.**
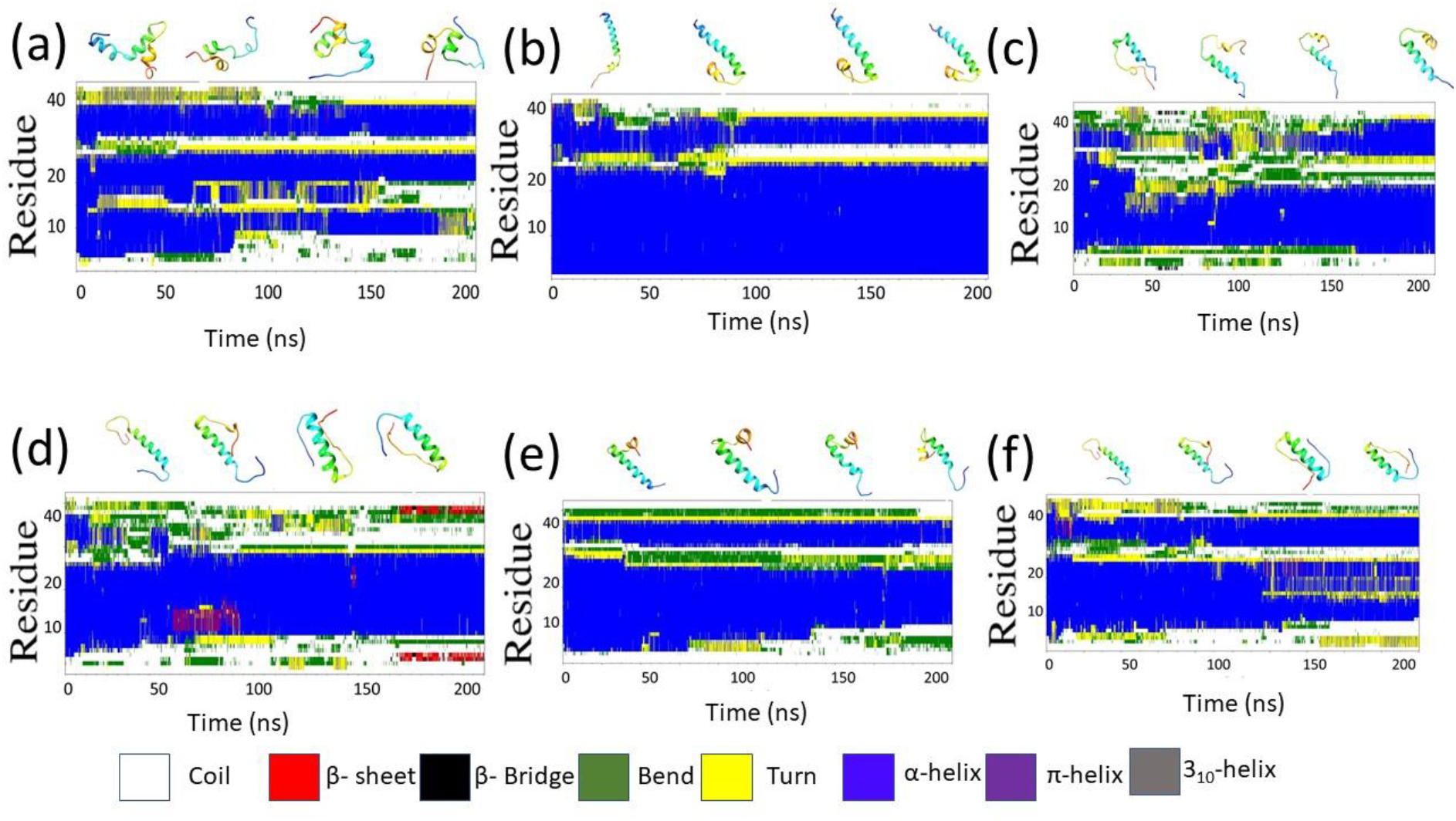
Time-dependent secondary structural changes of amyloid monomers and dimers on the surface of lipids and AQP4. (a) Shows the monomer at the surface of the POPC lipid bilayer, showing rapid transformation from an α-helix to a coil. (b) The presence of aquaporin-4 in the POPC membrane results in a slow transition and retains the α-helical structure. The α-helix is most stable and having less propensity for the aggregation. The amyloid monomers of dimers (c) and (d) show a transition from α-helix to coil and β-sheet formation. The inter-peptide interactions at the surface of POPC induces transformation to β-sheet structure with higher aggregation propensity. However, (e) and (f) show that AQP4 retains its α-helix in dimers. The aggregation of Aβ42 is associated with the transition from a α-helix conformation to a β-sheet-rich structure, whereas the α-helix conformation is less favorable for the aggregation.

**Figure 3.**
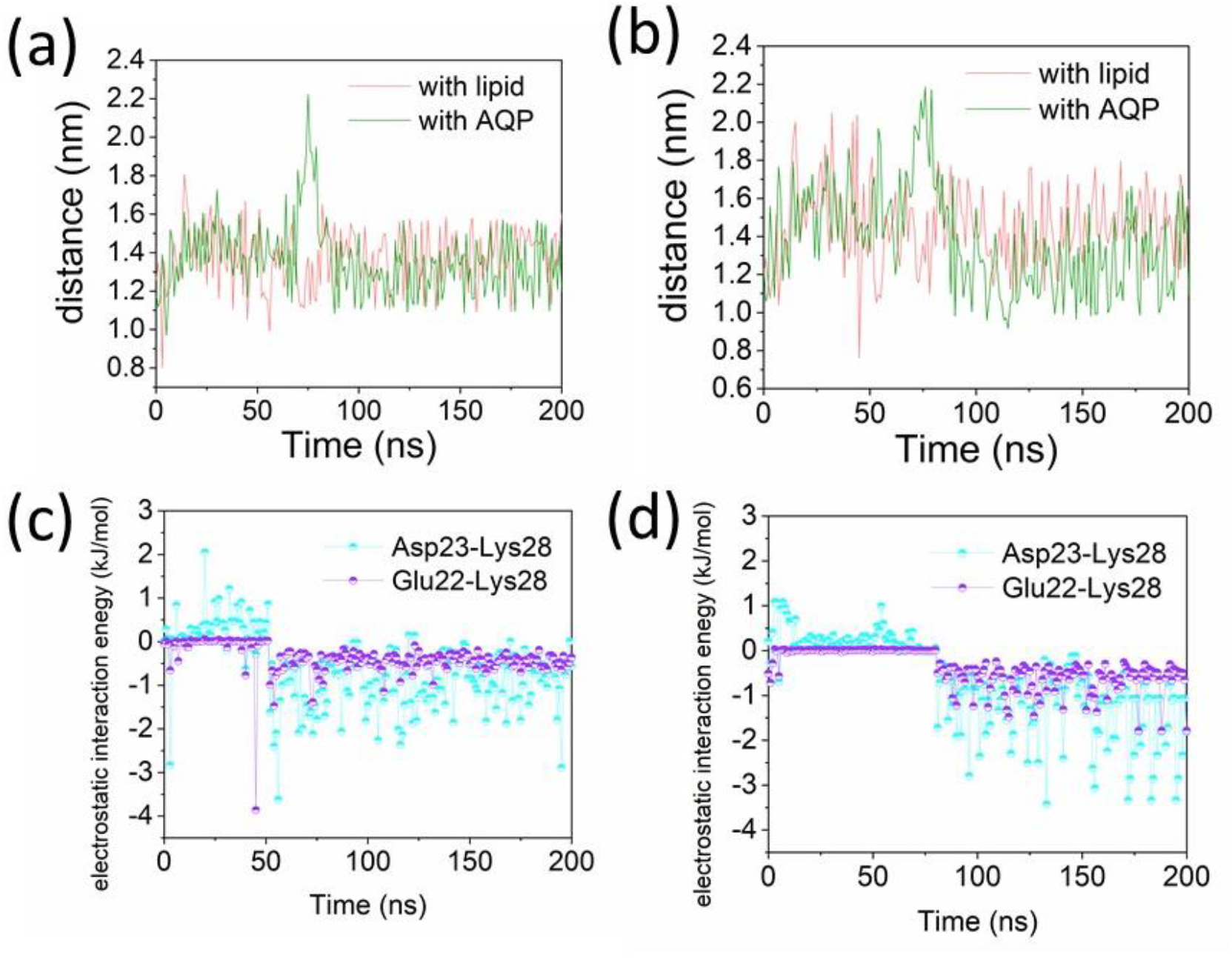
The salt bridge distances between Asp23-Lys28 and Glu22-Lys28 are essential because it stabiles the secondary structure of the peptide and reduces the aggregation propensity. In addition, it influences peptide interactions with surrounding molecules, including water. Salt bridge distances were calculated as the average distance between the COO^−^ and NH_3_^+^ moieties of the residues. (a) Represents the salt bridge distance between residues Asp23-Lys28, (b) represents the salt bridge distance between Glu22-Lys28. (c) and (d) represent the electrostatic interaction energies between residues Asp23-Lys28 and Glu22-Lys28 at the surface of POPC and AQP4.

**Figure 4.**
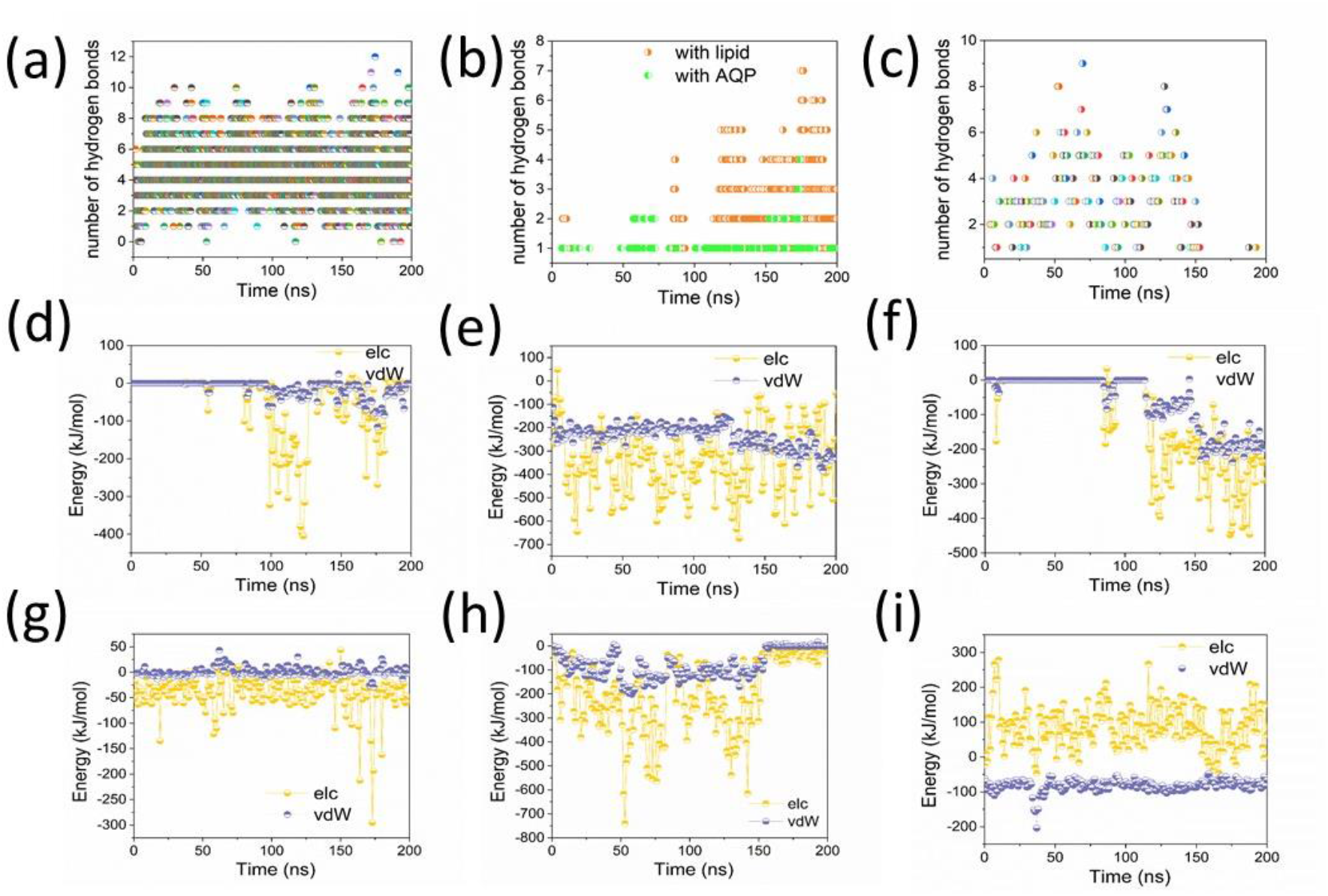
Hydrogen bond and non-bonded interaction energy analyses. (a) The number of hydrogen bonds between the amyloid monomer and AQP4. (b) Representative interpeptide hydrogen bonds in the dimer at the surface of POPC lipids and AQP4. Higher interpeptide hydrogen bonds indicate a higher propensity to aggregate the peptide at the lipid surface. (c) Number of hydrogen bonds formed between AQP4 and the entire dimeric peptide. (d) The electrostatic and van der Waals interaction energies between the monomer-POPC when AQP4 was present and (e) when AQP4 was not present in the membrane. (f) and (g) represent the electrostatic and van der Waals interaction energies of the interpeptides at the surfaces of POPC and AQP4, respectively. The individual peptide interaction energies of the dimers and AQP4 are represented in (h) and (i). Here, elc represents the electrostatic interaction energy and vdw represents the van der Waals interaction energy.

To understand the distribution of water around the amyloid peptide, a radial distribution function was estimated, as shown in Figure S1 of SI. The distribution function shows an identical pattern for both the POPC and AQP4 systems, but a higher shift (> 0.5 nm) was observed for POPC. The distribution of water around a peptide is influenced by the interaction of water molecules with AQP4 and conformational transition of the peptide. Furthermore, the hydrophobicity of the monomer reduced in the presence of aquaporin-4, whereas the hydrophilic area (Figure 5a-f) was stable. This is in agreement with the radial distribution function of monomers with water. Similarly, the individual residue solvent-accessible surface area demonstrated higher fluctuations in the presence of aquaporin-4 (Figure 6a-f). In addition, aquaporin-4 reduced the inter-peptide hydrogen bonds (Figure 4b) and increased hydrogen bonds with surrounding water (Figure S2 of SI). When the hydrophobicity of a peptide is reduced, the solvation of peptides is enhanced by providing a pathway for water molecules to interact with the hydrophilic regions. This enhances the hydrogen bonding with water, which can contribute to the stabilization of the peptides and reduce the propensity for aggregation. The electrostatic energy between water and the peptide was higher in the presence of AQP4 (Figure S3 of SI), indicating that the monomer units exhibited a strong interaction between the lipid headgroups at the surface and a higher degree of hydrophobicity. It retains its native α-helix structure due to the presence of AQP4 in the membrane, which draws the monomer towards the pore region through strong electrostatic interaction energy. A recent experiment involving amyloid peptides demonstrates that the formation of toxic oligomers is a significant factor in the onset of Alzheimer’s disease [45–47]. The dimer structure of the peptide on the lipid surface and the presence of aquaporin-4 were examined using all-atom MD simulations. The secondary structural changes in each monomer peptide of the dimer were independently assessed. The initial frame of monomer exhibited 48.4±1.6% helix, 51.6 ±1.6% coil and 51.8±0.5% helix, and 48.1±0.5% coil, respectively (Figure 2c and d). The interaction with aquaporin-4 and dimer restricted the secondary structural changes and preserved 52.4±0.1% helix, 47.6±0.1% coil, 75.5±1.5% helix, and 24.5±1.5% coil (Figure 2e&f) at the end of the simulation. This also suggests that aggregation depends on the inter-peptide interactions.

**Figure 5.**
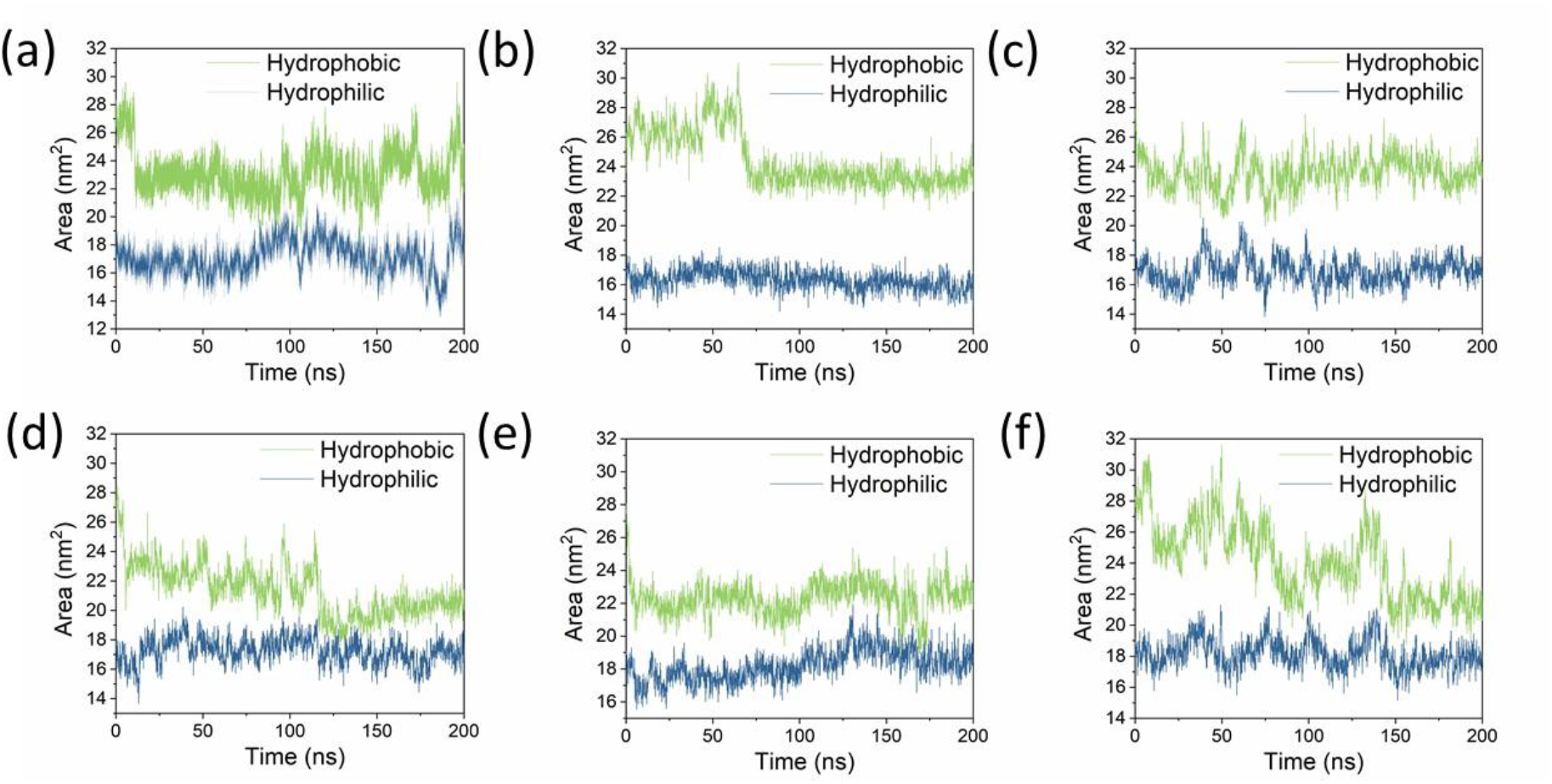
Hydrophobic and hydrophilic surface areas of the peptides. (a) represents the hydrophobic and hydrophilic areas of the amyloid monomer at the lipid surface and (b) represents the surface of AQP4. (c) and (e) represent the surface areas of one of the monomers at the surfaces of POPC and AQP4, respectively, and (d) and (f) represent the surface areas of the second monomer.

**Figure 6.**
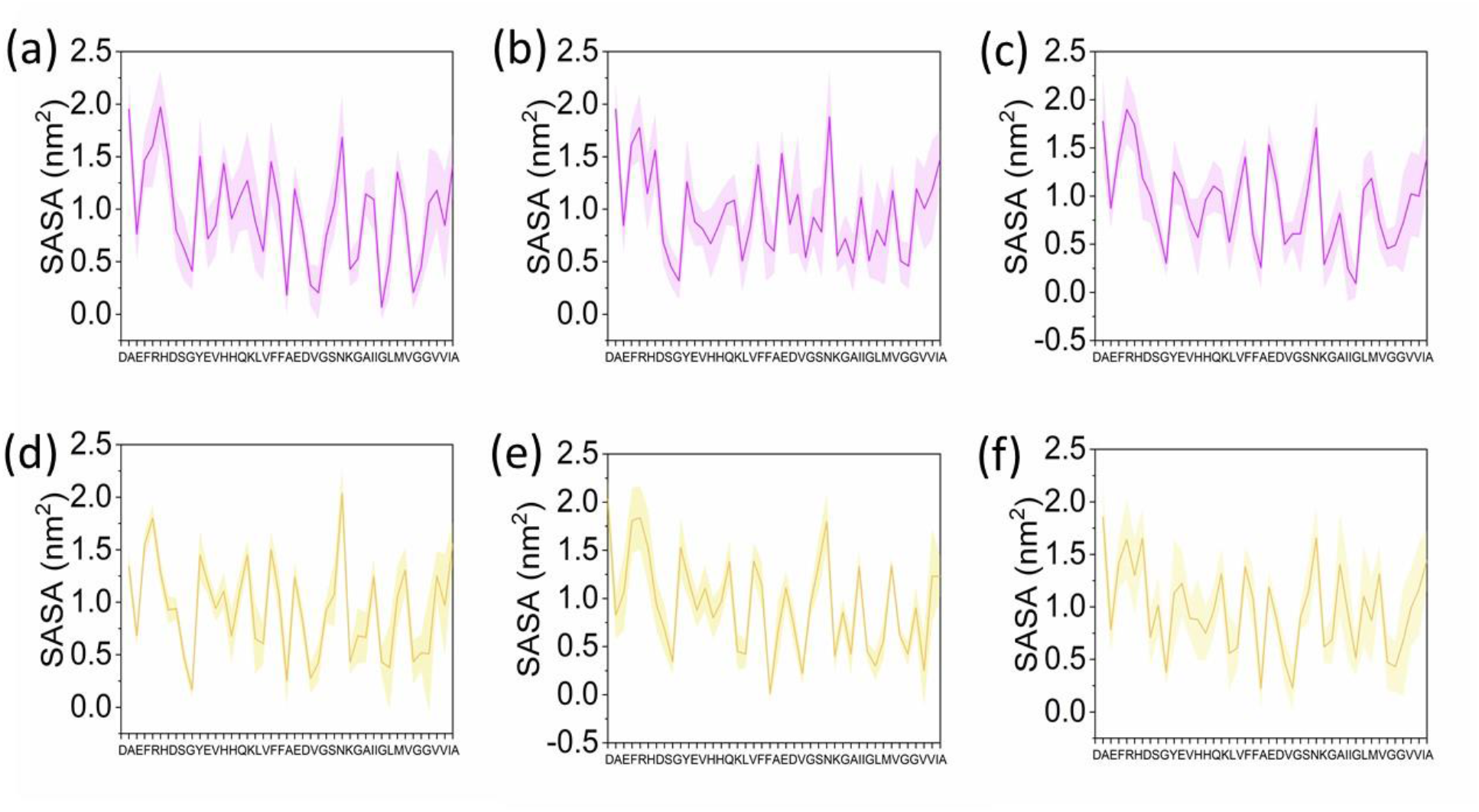
Individual residue solvent-accessible surface areas (SASA) of the peptide. (a)–(c) show the SASA residues of the monomers and monomers of the dimer on the surface of the POPC lipid bilayer. (d)–(f) represent SASA of the monomer and monomers of the dimer at the surface of AQP4.

### Aggregation at the surface of lipid and aquaporin-4 coarse-grained molecular dynamics simulation

The amyloid-β hypothesis implicates the ability of peptides to destabilize cell membranes, which ultimately leads to cell death [48–51]. Here, we used coarse-grained molecular dynamics simulations to understand the interactions among amyloid peptide oligomers, lipids, and aquaporin-4 within the cell membrane. By simulating these interactions over a long period (15 µs), we can gain a better understanding of how these interactions affect cell membrane function and how they may contribute to cell toxicity. Figure S4-S8 of Supporting Information illustrate the interaction of amyloid monomers with the surface of the lipid bilayer and at the surface of AQP4. The simulation results demonstrated that the amyloidogenic dimer interacts and aggregates at the surface of the lipid bilayer, in contrast to the amyloid monomer, which exhibits attraction towards AQP4 and fewer interactions with the lipid bilayer. Nonetheless, when moving from the trimer to the pentamer, aggregation occurs in water and there are occasional interactions with the lipid bilayer over the 15µs. In contrast, oligomers (trimers, tetramers, and pentameters) are attracted to AQP4 prior to aggregation in an aqueous environment. This suggests that the presence of AQP4 at the surface of the lipid bilayer hinders aggregation, resulting in strong attraction towards AQP4. By analyzing the deuterium order parameter, lipid density, lipid surface area, and lipid thickness, we assessed lipid stability during the all-atom simulation (Figure 7a-c). A higher-order parameter was observed in aquaporin-4 embedded lipid membranes. This is due to the presence of transmembrane proteins that stabilized the membrane through hydrophobic matching and electrostatic interactions. Moreover, membrane thickness was stable in the presence of aquaporin, whereas simulations of the lipid system revealed abrupt fluctuations in membrane thickness. This demonstrates the potential toxicity of Aβ to lipid membranes. Variations in membrane thickness and order parameters suggest that amyloid-β exerts detergent effects on the membrane. Owing to the temporary and dynamic equilibrium between dimers, trimers, and tetramers, it is difficult to define early oligomers at the atomic level using biophysical techniques. Early oligomers were the most hazardous species throughout the aggregation process. The formation of low-molecular-weight oligomers from a combination of dimers, trimers, and tetramers is characteristic of amyloid fibrils. Several simulation studies have investigated the free energy surface and dynamics of dimers, which serve as building blocks for larger oligomers [52–55] also the cell membrane is the primary target of amyloidogenic oligomers [56–57]. The amphipathic nature of amyloid peptides may facilitate their interactions with the membrane surface and insertion into membrane cells. In addition, experimental investigations have indicated that membrane interactions promote amyloid fibril formation [58–62]. Using all-atom simulations, we showed the formation of a β-structure in the dimer peptide upon interaction with lipid bilayers, although these modifications were inhibited in the presence of AQP4. Furthermore, the presence of AQP4 in the membrane resulted in a significant reduction in peptide-peptide interactions. The interaction between the peptides and lipids disrupts the membrane structure, resulting in dysregulation of ion and water homeostasis [63–67]. Even though there are no particular binding sites, the interaction of the amyloidogenic peptide with the membrane surface results in asymmetric pressure between the two lipid bilayers, uncontrolled leakage of small molecules, and detergent-like effects [66]. Peptide aggregation is dependent on various factors, including the membrane type, pH, peptides, and salt concentration [68]. A negatively charged membrane is known to produce a β-strand structure, which further perturbs the membrane structure [69]. The aggregation of peptides is also dependent on their charge and orientation; the peptide-peptide interactions rely on the orientation of the peptides in the environment, which results in the formation of fibrils on the surface of the membrane. As observed in the snapshot (Figure S4), the monomers interface with the membrane by electrostatic interactions, and the peptides align parallel to the membrane, whereas the oligomers form peptide-peptide interactions and aggregate at the membrane surface. To reveal the structural alterations, the structure of the peptide from the last frame of CGMD was retrieved and transformed into an all-atom model. In the presence of AQP4, post-CG all-atom simulations revealed a higher helical content. At the lipid surface, the secondary structure of the peptide showed 12.8% helices, 41.0% turns, and 46.2% coils, whereas for AQP4, 33.3% helices, 20.5% turns, and 46.2% coils. The salt bridge obtained from post-CGMD between Asp23-Lys28 and Glu22-Lys28 was consistent with the all-atom simulation. This demonstrated the retention of helical content of the peptides while interacting with AQP4, and the transformation of peptides into a more toxic β-sheet structure when interacting with the lipid bilayer. Salt bridges help stabilize the helical structure of the peptide, and the positioning of these residues helps form strong hydrogen bonds that stabilize the α-helix. When salt bridges are broken, the peptide is transformed into a more toxic β-sheet. β-sheets are more toxic because of their high propensity to form insoluble aggregates, which leads to the formation of amyloid plaques. Due to the salt bridges, the strong hydrogen bonds between the residues do not show interpeptide interactions that lead to aggregation.

**Figure 7.**
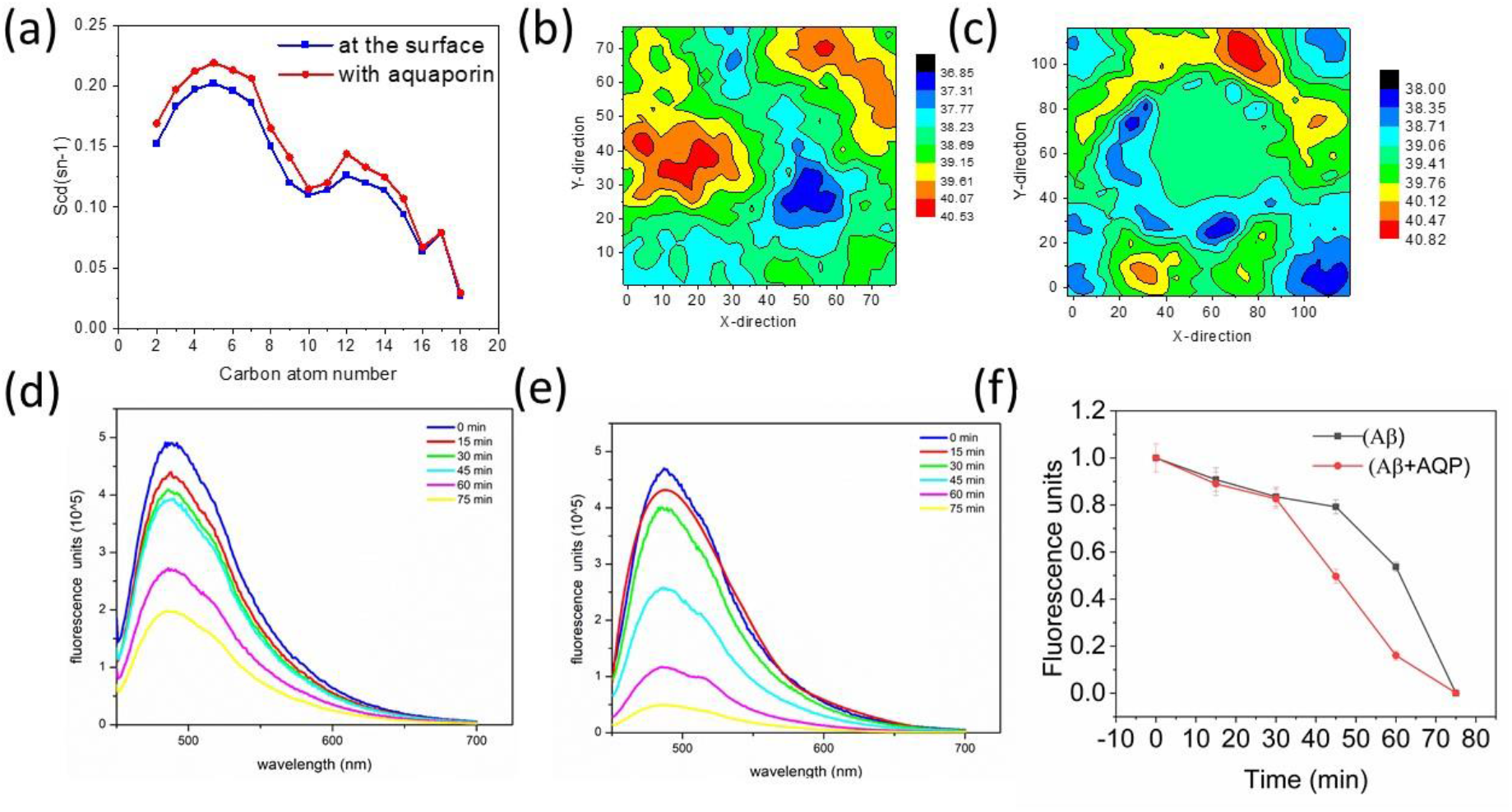
Peptide aggregation and lipid stability. (a) The lipid order parameter obtained for the peptide at the surface is represented in blue, and the peptide with aquaporin embedded in the membrane is represented in red. The higher lipid order for the aquaporin-embedded membrane indicates higher stability of the membrane. Lipid thickness from all-atom simulation of peptides at the surface of (b) POPC and (c) AQP4. (d)–(f) represent thioflavin T emission fluorescence spectra (range 450-700 nm) with excitation at 446 nm for the peptide and the peptide with AQP4 in vitro. A Higher Thioflavin T emission intensity indicates a higher extent of aggregation. The peak was higher in the absence of AQP4, as indicated in (d). The presence of aquaporin resulted in reduced intensity, which was correlated with the slower aggregation of the peptide. The normalized time-dependent aggregation of peptides (f) indicates a higher intensity for peptides than for peptides with AQP4, indicating that aquaporin reduces peptide aggregation.

### Time-dependent aggregation of amyloid peptide-Thioflavin fluorescence spectroscopy

In *vitro* amyloid-β aggregation was evaluated using thioflavin-based fluorescence emission. Thioflavin-T, a yellow fluorescent dye, binds to amyloid fibrils and produces an intense specific emission at 490 nm. Therefore, we used this assay at varying times and concentrations to monitor the AQP4-mediated aggregation of amyloid peptides. Here, we adapted an optimized procedure by Barnoli et al. to perform this assay [70]. The aggregation process of the peptide was time dependent, and aquaporin-mediated aggregation was slower than that of amyloid-β alone under the same conditions. Amyloid-β can adopt two different conformations in solution such as anti-parallel β-sheet and random coil or α-helix structures. The former is highly amyloidogenic, and the latter is poorly amyloidogenic. The time-dependent aggregation based on fluorescent intensities is depicted in Figure 7d-f. After lyophilization, the amyloid-β from the HFIP solution was dissolved in DMSO. Aliquots of 2 µL amyloid-β in dimethyl sulfoxide (DMSO) were incubated alone or with AQP4 at a molar ratio of 100:1 in 0.215 M sodium phosphate buffer (pH 8.0). The aggregation of the peptide occurred rapidly, and the intensity was measured every 15 min. The incubated aliquots were selected based on 15-minute intervals up to the lowest peak obtained (at 75 min). Several studies have reported that amyloid-β aggregates form cross-β fibrils *in vitro,* similar to that observed in patients with AD. The fluorescence intensity of amyloid-β was found to be higher than amyloid-β+AQP4 from 0-75 min. The presence of AQP4 shows slower aggregations as the intensity is found to be lower than amyloid-β (Figure S9 of SI). It was also observed that higher concentrations of AQP4 resulted in slower aggregation *in vitro* (Figure S10 of SI).

### Water dynamics of aquaporin-4

Figure 4d-i depicts the van der Waals and electrostatic interaction energies between the amyloid fragments and AQP4. The interaction energy between the amyloid monomer and lipids was stronger in the absence of AQP4. An average of -190 kJ/mol van der Waals and -150 kJ/mol electrostatic interaction energies were observed between the monomer peptide and POPC, respectively. However, the presence of AQP4 at the membrane reduced the energies to -3 kJ/mol and -45 kJ/mol van der Waals and electrostatic interaction energy respectively. In addition, the monomeric peptide with AQP4 showed van der Waals and electrostatic interaction energies of -217 kJ/mol and -398 kJ/mol, respectively. This strong electrostatic interaction indicates that AQP4 attracts amyloid fragments towards the pore region. In addition, the interpeptide hydrogen bonds in the dimer were stronger with lipids (2-8 nos) than with AQP4 (1-3 hydrogen bonds). However, 12 hydrogen bonds were observed between the monomer peptides and AQP4, indicating a stronger affinity of the peptide for AQP4. The interaction between the peptides and AQP4 results in structural modifications of the pore region and helix bundles located near extracellular regions. The pore region of aquaporin-4 is primarily controlled by residues Arg216, Ala210, Phe77, and His201 [71]. These residues are positioned opposite to each other and regulate pore diameter for water molecule transport. To understand the dynamics of the pore and its associated residues, we calculated the distance between residues Arg216-His201 and Ala210-Phe77 (Figure 8a&b). AQP4 interacts with the amyloid peptide monomer and causes a shorter distance between Arg216 and His201. However, the dimer interactions resulted in the highest distance of 0.9 nm. Additionally, we observed that the interaction of the peptide with the pores due to steric hindrance led to structural changes and resulted in higher distance between residues Phe77 and Ala210. Furthermore, peptide interactions not only induced structural modifications but also altered the hydrogen bonding pattern between the critical residues and water (Figures 9b and c). Peptide interactions also led to higher hydrogen bonds between Arg216, Ph77, Ala210, and His201 and water molecules. This is due to the large number of water molecules occupying the pore region, as depicted in Figures 9b and c and 10b and c. In addition, the interaction of these residues with peptides disrupted their natural orientation in the pores (Figure 8 h and i). The constricted pore entrance widened, allowing more water molecules to pass through the AQP4. The helical structure of AQP4 is important because it affects the structural stability and confined size of the pores in the region containing NPA motifs. As depicted in Figure S11-S13 of the SI, time-dependent secondary structure analysis revealed stable helical structures of AQP4 at POPC(h1-h8). However, the interaction of peptides generates occasional coil, bend, and turn formations in the h7 helices. Additionally, the α-helices of the h8 region exhibit transitions to coil and bend/turn. H2 helices contain Phe77 residues, and the coil-to-turn transition found in secondary structural alterations implies relocation of residues near the pore region (Figure 8 h &i). In addition, the transition observed in the small helices pushes the residues to the pore region and collapses the AQP4 cavities, leading to the entry of several water molecules rather than a single-line flow (Figure 9 b&c).

**Figure 8.**
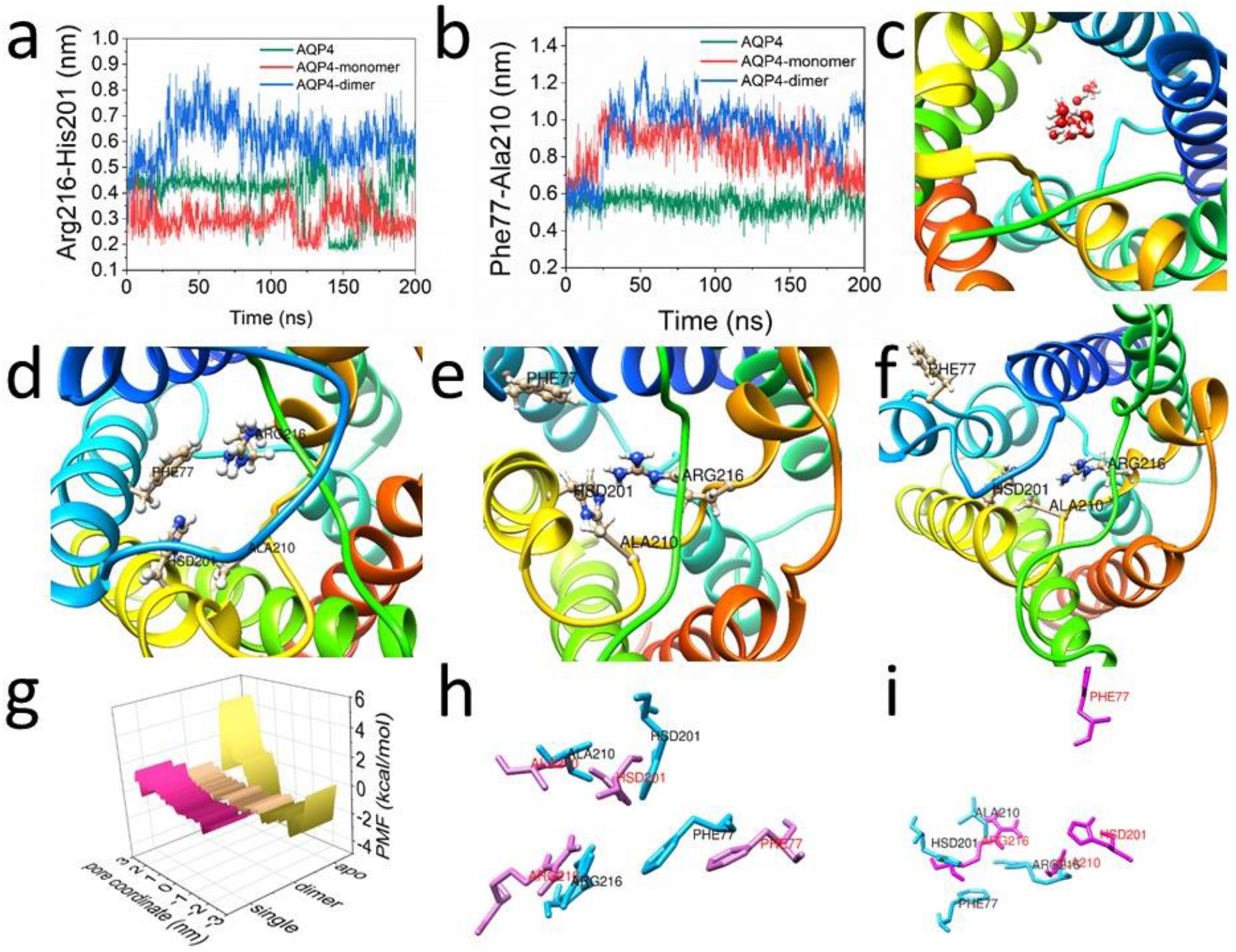
The dynamics of AQP4. (a) and (b) represent the distances between residues in the pore region of AQP4. These residues control the pore diameter and selectivity of water molecules through the channel. Interactions between peptides were affected by inter-residue distances, resulting in pore widening. (c) Vertical view of normal single-line flow of water molecules. (d) Vertical view of residues Arg216, His201, Phe77, and Ala210 in the pore region of native AQP4; (e) and (f) represent the same when interacting with the amyloid monomer and dimer. The energetics of water flow through the channel were calculated by obtaining the PMF profile (g), which showed a high barrier at the selectivity filter and was broken upon interaction of the monomer and dimer peptides. (h) and (i) represent the alignment of the residues in the pore region while interacting with the monomer and dimer peptides, respectively.

**Figure 9.**
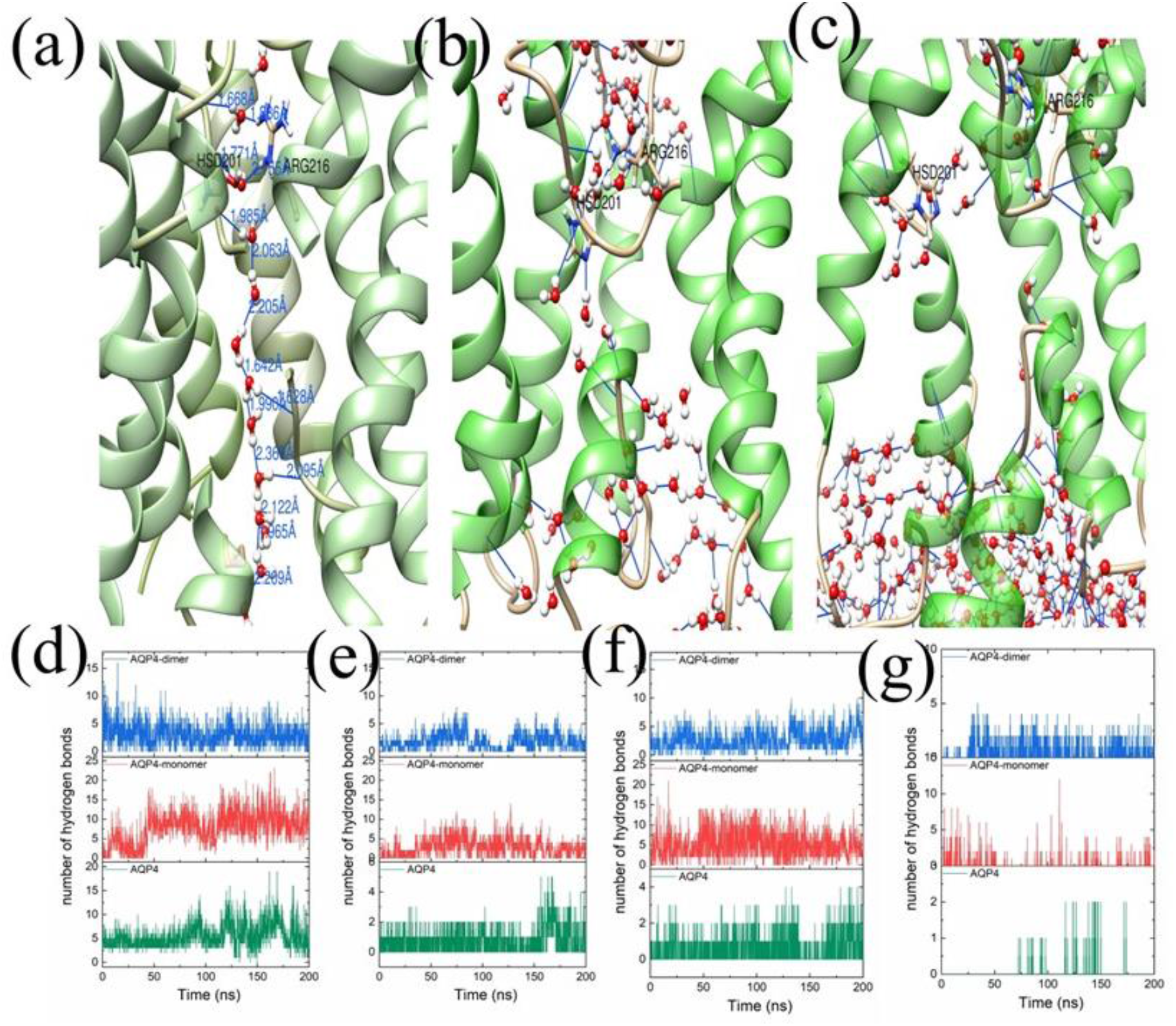
The single-line flow of water through the channel was achieved by creating and breaking hydrogen bonds. (a) represents the single-line flow of the AQP4 channel, and (b) and (c) represent the disrupted water flow through the channel while interacting with amyloid monomer and dimer peptides, respectively. (d)–(g) represents the number of hydrogen bonds formed between water molecules and residues Arg216, Ala210, His201, and Phe77, respectively.

To elucidate the peptide-induced changes in AQP4, water permeation through both heteromeric and monomeric aquaporin-4 was assessed. Using the single-AQP4 diffusive permeability constant and osmotic water permeability constant, water permeation through AQP4 was determined. Along the +z and − z axes, Aquaporin-4 monomer units showed a total of 172 and 175 water permeation events, respectively. Under normal conditions, a single unidirectional flow of water molecules across aquaporin-4 and AQP2 has been reported in earlier studies. The interaction of amyloid peptides disrupts AQP4 structure and water flow. The interaction of monomer peptides with AQP4 resulted in a net flow of 8396 water molecules, significantly increasing the water flow through the native AQP4 monomer, which initially contained only 836 water molecules. Additionally, the interaction of AQP4 with dimeric peptides further increased the water flow to 16372. The calculated net water flow and permeability coefficients are presented in Table 1. The interaction of the amyloid-β peptide with the pore region interrupts the single-line flow of water, and water was shown to occupy regions of the α-helix other than the central pore. As shown in Figure 10a, eight water molecules form a single file in AQP4. Nevertheless, the peptide-AQP4 interaction resulted in a bulk-like flow of water, as depicted in Figures 10b and c. This may result in loss of the normal hydrogen-bonding pattern and probable ion or proton permeation. In addition, the estimated potential of the mean force (PMF) along the direction of a single water movement revealed conformational changes induced by the peptides. At the selectivity filter, the native AQP4 has the greatest barrier height of 4.85 kcal/mol, but the AQP4 that interacted with single and dimer peptides have barrier heights of 1.74 kcal/mol and 0.59 kcal/mol, respectively. It was also noticed that the PMF of the monomer and dimer interactions showed a collapsed selectivity filter, and water molecules passed through AQP4 indiscriminately (Figure 8g). Figure 10d-f demonstrate a disturbance in the regular functioning of AQP4 and significant changes in its diameter, as indicated by the HOLE radius. This further implicates that the physiological abnormalities of the brain reported in patients with Alzheimer’s disease are related to amyloid accumulation. The brains of Alzheimer’s disease patients exhibit a deterioration of normal tissue architecture and an edematous or ’dry’ characteristic. Vascular damage and reactive gliosis were also found in the majority of patients with AD, accompanied by ischemic astroglial cells and APP protein [72–73]. Several experimental studies have demonstrated the significance of AQP4 in the etiology of Alzheimer’s disease and clearance of β-peptide [74–79]. However, the mechanism through which Aβ is eliminated remains unclear. Molecular dynamics simulations demonstrated that the peptides were unable to pass through AQP4. Previous studies have revealed a correlation between altered astrocyte function and Alzheimer’s disease, with the presence of AQP4 in astrocytes influencing calcium and potassium homeostasis. [16,80]. According to PMF calculations, the decreased height of the barrier implies the loss of normal AQP4 function: the transport of water and blocking of ions, including H^+^, are disrupted. Larger ions such as K^+^ and Ca^2+^ can pass through lower barriers and wider pores [14,16,]. AQP4 knockout animals exhibit an increase in Aβ deposition in the cerebral cortex of 5xFAD mice; conversely, inducing the expression of AQP4 decreases Aβ levels in brain tissues [81–82]. In addition, the presence of water in the CSF is altered in transgenic mice with amyloid deposits owing to the loss of native AQP4 function.

**Figure 10.**
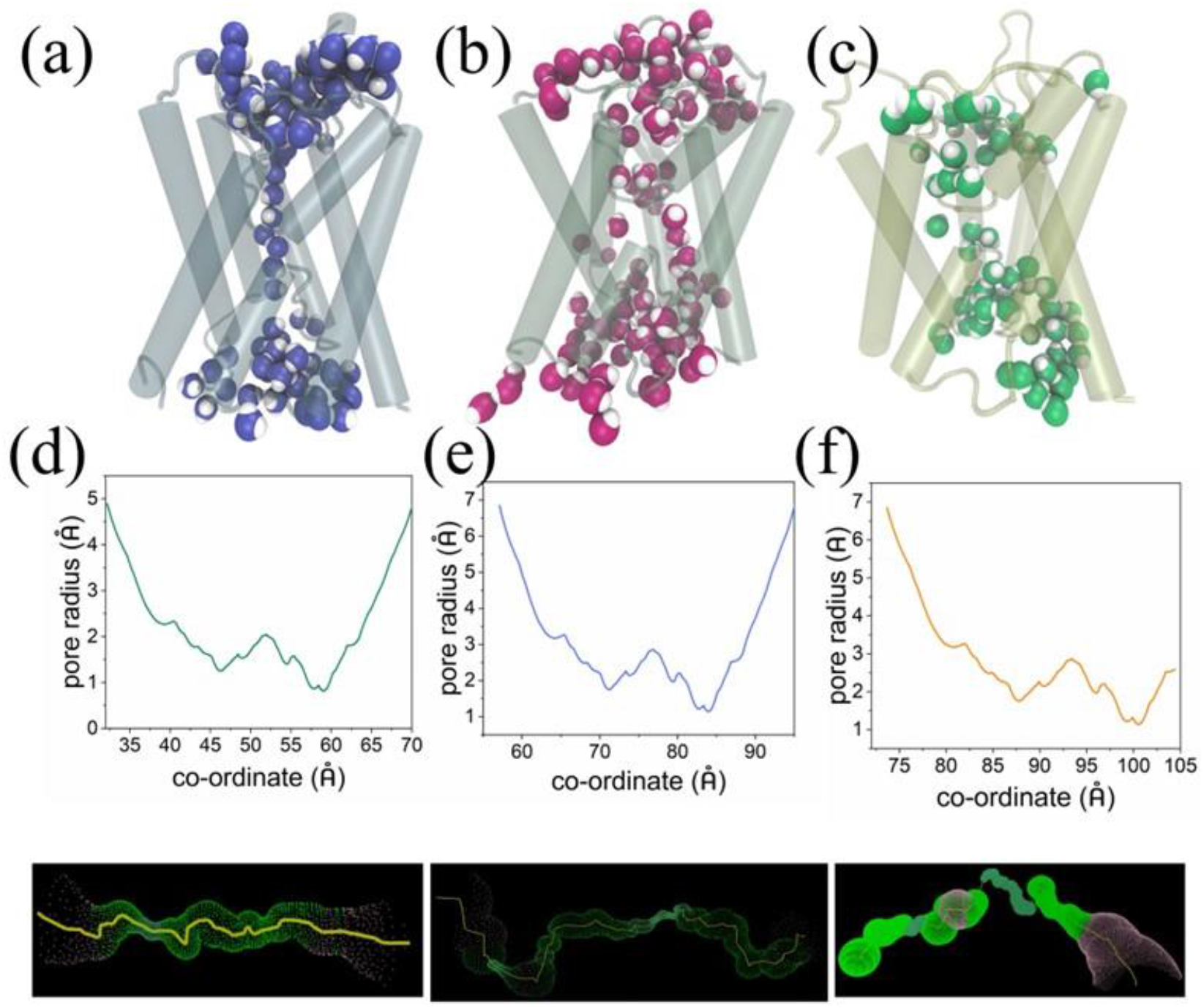
Single-line flow of water and pore structure. (a) represents the aquaporin-4 single-line flow of water. (b) and (c) represent the disrupted water flow owing to the interaction of the monomer and dimer peptides, respectively. The interaction between amyloid monomer (b) and dimer (c) leads to bulk water flow through the channel. The pore diameters (d-f) also indicate loss of the selectivity filter in the channel.

**Table 1.** Permeability coefficients calculated from equilibrium MD simulations.

| | $P_d$<br>($10^{-14} \text{ cm}^3/\text{s}$) | $P_f$<br>($10^{-14} \text{ cm}^3/\text{s}$) | The net flow of<br>water molecules<br>along the z-direction |
| --- | --- | --- | --- |
| AQP4 | $2.42 \pm 0.2$ | $5.08 \pm 0.5$ | 836 |
| AQP4 with<br>monomer | $6.2 \pm 0.5$ | $8.6 \pm 0.7$ | 8396 |
| AQP4 with dimer | $12 \pm 0.1$ | $15.51 \pm 0.6$ | 16372 |

### Protein-Protein interaction network from text mining

Analysis of the protein-protein interaction network revealed the co-occurrence of Aquaporin-4 with amyloid-β. User-specific keywords were utilized to retrieve scientific literature from the PubMed database, and our earlier approaches were modified to identify the gene and protein entities mentioned in the literature. A database dictionary was developed for proteins/genes with specific goals, including Aquaporin-4, amyloid-peptide, Alzheimer’s disease, and amyloid precursor proteins. The biological relationship between these established co-occurring proteins and the discovered proteins/genes is depicted in Figure S14 of the SI. Employing the identified proteins/genes and their relationships, Gephi 0.10 was used to construct the interaction network. In addition, several network analysis algorithms, including centrality measures, page rank, node degree, and sub-net-node degree analysis, were applied to the network to identify gene/protein hub and dense protein clusters. The network demonstrated that AQP4 and APP are interconnected, along with several Alzheimer’s disease-related proteins (Figure S12). The extensive network diagram with all the connected proteins illustrated the numerous proteins involved in Alzheimer’s disease, as well as apoptosis-related proteins such as MAPK, BCL2, and CASP1. A previous report revealed the co-occurrence of AQP4 proteins and apoptosis induced by T-2 toxin-induced neuronal stress [71]. The extensive network contained 1140 nodes (proteins) and 2276 edges (interactions). Network analysis led to the identification of several proteins and linked pathways connected with AQP4-mediated amyloid peptide aggregate formation and Alzheimer’s disease. In addition, we identified 13 pathways associated with Alzheimer’s and AQP4, including axon guidance signalling, lysosome signalling, hedgehog signalling, Notch signalling, apoptosis, proteasome, MAPK signalling, pi3kakt signalling, insulin signalling, toll-like receptor signalling, pentos phosphate signalling, calcium signalling, and CAMP signalling pathways. The involvement of these pathways is either directly or indirectly related to the expression of AQP4 in the brain and clearance of Aβ, and extensive experimental studies are needed to reveal the relationship and role of these pathways in the clearance of Aβ. Identifying co-occurring genes or proteins can provide significant insights into the relationships and interactions between different biological entities. This information helps us to understand the “missing” biological pathways and their interplay during the clearance of Aβ. In addition, from the extensive network depicted in the network diagram, we identified key proteins related to AQP4 and Alzheimer’s disease. These proteins are also recognized in a variety of biological processes that may be linked to Alzheimer’s disease and AQP4.

In conclusion, our study established a significant correlation between AQP4 expression and amyloid aggregation. We used text mining to construct a network of protein-protein interactions and revealed that AQP4 plays a crucial role in the etiology of Alzheimer’s diseases by interacting with several interconnected proteins and pathways. Our study revealed the nature of aggregation at the surface of the POPC lipid bilayer and its interaction with amyloid oligomers. Peptides interact with the pore region of AQP4 and disturb the orientation of critical residues. However, this interaction maintained the helical content of the peptide while reducing its toxicity at the lipid surface. Furthermore, the interaction also reduced the hydrophobicity of the peptide, thereby accelerating its removal from the astrocytes. Long-term coarse-grained MD simulations demonstrated different features of oligomer aggregation at the surface and substantial oligomer attraction to aquaporin, which inhibits the aggregation process. Moreover, we observed that the selectivity filter for AQP4 was destroyed, thereby disrupting water flow. Our findings also shed light on the physiological changes in brain tissue associated with Alzheimer’s disease, including water retention and increased water flow in the CSF. Finally, we demonstrated aquaporin-4’s ability to minimize aggregation over time, using in vitro thioflavin fluorescence spectroscopy.

## Materials and methods

### Protein-Membrane simulation system

The X-ray crystal structure of AQP4 [83] and the NMR structure of Alzheimer’s amyloid-peptide [84] were obtained from the PDB databank using PDB IDs 3GD8 and 1IYT, respectively. The AQP4 tetramer was embedded in the (1-palmitoyl-2-oleoyl-sn-glycero-3-phosphocholine) POPC lipid bilayer after removing all co-occurring molecules, including water. Before membrane insertion, the OPM web server [85] was used to align proteins with normal membranes. A POPC membrane bilayer with 120 Å lengths in the x- and y-directions was constructed using the CHARMM-GUI web server [86]. To maintain a real simulation system, a 30 Å thick water box was added below and above the protein membrane complex, and the system was neutralized with KCl ions.

### All-atom molecular dynamics simulation

All-atom molecular dynamics simulations of the protein-membrane system were performed using the GROMACS 2019.6 [87–88] simulation package with the CHARMM36 forcefield [89] and TIP3P water model [90]. The particle mesh Ewald method [91] was used for the calculation of electrostatic interactions, and the Verlet cut-off scheme with a distance of 1.4 nm was used for short-range repulsive and attractive interactions. All the bonds were contained using LINCS algorithms [92], and Nose-Hoover temperature coupling along with the Parrinello-Rahman algorithm was used to maintain the temperature and pressure at 310 K and 1 bar, respectively. The simulation was performed for 200 ns using the NPT ensemble. The minimization of the system was performed using the leapfrog algorithm, followed by equilibration using the NVT and NPT ensembles for 20 ns. The trajectories were collected every 100 ps, and the periodic boundary effects were corrected before analysis. The simulation results were validated by repeating the simulation with different initial velocities. The water dynamics of the aquaporin channel were evaluated by calculating the number of water molecules present in the top, bottom, and channel. The final data represent the mean of the total water molecules crossing the channel. The water permeability was calculated from the equilibrium MD simulation [93] using the following equations

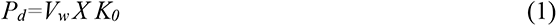

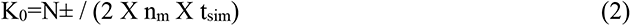

Here, V_w_ represents the average volume of a single water molecule and k_0_ represents the number of water molecules that cross per unit of time. Nm represents the number of monomer units, and t_sim_ is the total duration of the simulation. Furthermore, the permeability coefficient (pf) calculated from the collective diffusion constant, defined according to the Einstein relation as follows:

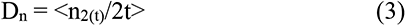

Where <n2(t)/2t> is the mean square displacement of n(t), the osmotic permeability constant (pf) calculated as

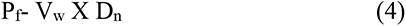

### Steered molecular dynamics and umbrella sampling

The free energy for the transportation of water molecules through the AQP4 channel was obtained by constructing the potential of mean force (PMF). The PMF profiles provide the energy required for a water molecule from one end of the channel to the other. Here, we extracted the last frame of AQP4 from 200 ns all-atom simulation. The PMF was calculated for AQP4 and AQP4 that interacts with amyloid-β peptides. The water molecules were placed ∼5 Å away from the pores of the channel and pulled through the channel cavity. The simulation box was maintained to satisfy the minimum image convention and provide sufficient space for pulling along the z-axis. The system was equilibrated with an NPT ensemble for a short period of 1 ns and minimized for 5000 steps. The temperature and pressure were maintained at 310 K and 1 bar, respectively, using the Berendsen weak coupling method. Pulling simulations (SMD) were performed by keeping the channel as a reference group with a harmonic spring constant of 900 kJmol-1 nm-2 and pulling rate of 0.001 nmps-1. The histograms were generated with a window spacing of 0.2 nm and approximately 450 frames corresponding to the COM distance between the reference and pulling groups. Each window was used to perform a 10 ns umbrella sampling simulation, and the PMF profile was constructed for each molecule, which provided the ΔG value for the transportation of water molecules. The weighted histogram analysis method (WHAM) was used to estimate the free energy profile from umbrella simulations [94].

### Coarse-Grained molecular dynamics simulation

Simulation system of Aquaporin-4 embedded lipid bilayer with amyloid-β was constructed with a Martini builder with the help of the CHARMM-GUI web server. In the Martin model, heavy atoms such as C, N, and O and their associated hydrogen atoms are considered as single beads. The simulation of 15 µs was performed with GROMACS using standard martini v2.2 forcefield [95–96] for proteins, lipids, water, and ions. The steepest descent algorithm was used to minimize the neutralized system, and equilibration was achieved with multiple steps by adapting the stepwise release of restraints applied to lipids. The periodic boundary conditions were applied and a time step of 30 fs was used electrostatic interaction was truncated with a shift cutoff of 1.2 nm and Lennard-Jones (LJ) potential was smoothly shifted to zero within 0.9 to 1.2 nm during the simulations. The temperature and pressure were maintained at 310 K and 1 bar using the Berendsen and velocity-rescaling methods. The trajectories were saved every 20 fs, and all visualizations were performed using VMD software packages [97]. The HOLE program [98] was used to calculate the HOLE radius of the AQP4. HOLE radius provides the size and shape of the AQP4 pore and cavity.

### Protein-Protein network analysis from text mining

A protein-protein interaction network was constructed to understand the protein associated with AQP4 and amyloid β-peptide, including Alzheimer’s disease. Here, we used our previous method [99,74], which harvests entries from PubMed using suitable keywords. The keywords such as “AQP4” and “aquaporin-4” returned a total of 4820 documents and amyloid β-peptide including Alzheimer’s disease returned 415,321 entries, respectively. The extracted data were used to construct a relation network using Gephi 0.10 [100]. Several network analysis algorithms and parameters, such as centrality measures, page rank, node degree, graph density, and sub-network analysis, were performed on the network. Centrality measures provide highly dense or the most important highly connected nodes in the network. The pagerank algorithm determines the relative importance of a particular node in a network based on its network connectivity. Furthermore, we reduced the broad network obtained from the above keywords into a small network where the most important proteins were identified.

### Thioflavin T-based fluorometric assay

Thioflavin T, amyloid-β peptide, DMSO, HFIP, and other reagents were purchased from Sigma-Aldrich. The aquaporin-4 and amyloid-β antibodies and proteins were obtained from ElabScience. A stock solution of Aβ_1-42_ (2 mg/ml) was prepared by suspending 1 mg of Aβ_1-42_ with 0.5 mL of 1,1,1,3,3,3-hexafluoro-2-propanol (HFIP). Rapid aggregation of Aβ_1-42_ was prevented by water batch sonication for 10 min. Aliquots of the peptide were stored at -80 °C, and freshly thawed samples were used for analysis. The thawed samples were sonicated until the HFIP evaporated. Further, sodium hydroxide solution was used to titrate the solution to a pH range of 8-9. For the co-incubation experiments, a buffer solution containing both the peptide and aquaporin was added at a molar ratio of 100:1 (AQP4: Aβ_1-42_) to conduct the thioflavin T fluorescence experiments [70]. Incubation was performed for a duration of 72 hours. The thioflavin T-based fluorometric assay was performed with Horiba Jobin–Yvon Fluoromax 4 using a 3 mL quartz cell. Fluorescence was monitored at 446 nm and 490 nm with excitation and emission slits of 2 nm bandwidth, respectively. The fluorescence emission spectrum was recorded between 450 and 600 nm, with excitation at 446 nm. The sample was analyzed after incubation, and the solution containing Aβ_1-42_ and Aβ_1-42_ with aquaporin-4 was added to 50 mM glycine-NaOH buffer (pH 8.5) containing 1.5 µM thioflavin T in a final volume of 2 ml. A time scan of the fluorescence was performed, and the intensity values reaching the plateau were averaged after subtracting the background fluorescence from thioflavin T and AQP4.

## Associated content

### Supporting Information

The supporting information contains a radial distribution function, representation of a coarse-grained simulation snapshot, secondary structural changes of AQP helices, and network diagrams.

## Acknowledgement

The author is thankful to Prof. Ponamali Kolandaivel for their support and encouragement.

## Author Contribution

NM designed the work, performed the simulations and experiments and prepared the manuscript.

---

**Figure S1.**
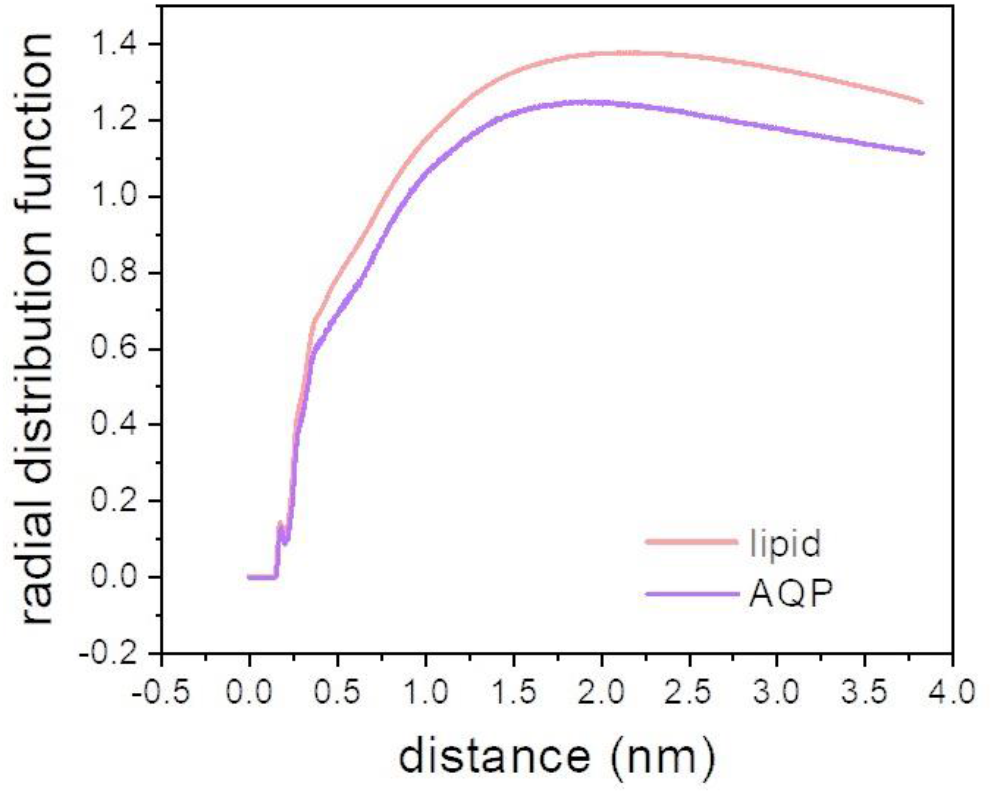
Radial distribution function of the amyloid monomer with water at the surface of POPC and AQP4. The differences in the RDF values suggest that the dynamics or interaction of water with peptides is influenced when AQP4 is present. Water is less dynamic with respect to peptides when AQP4 is present.

**Figure S2.**
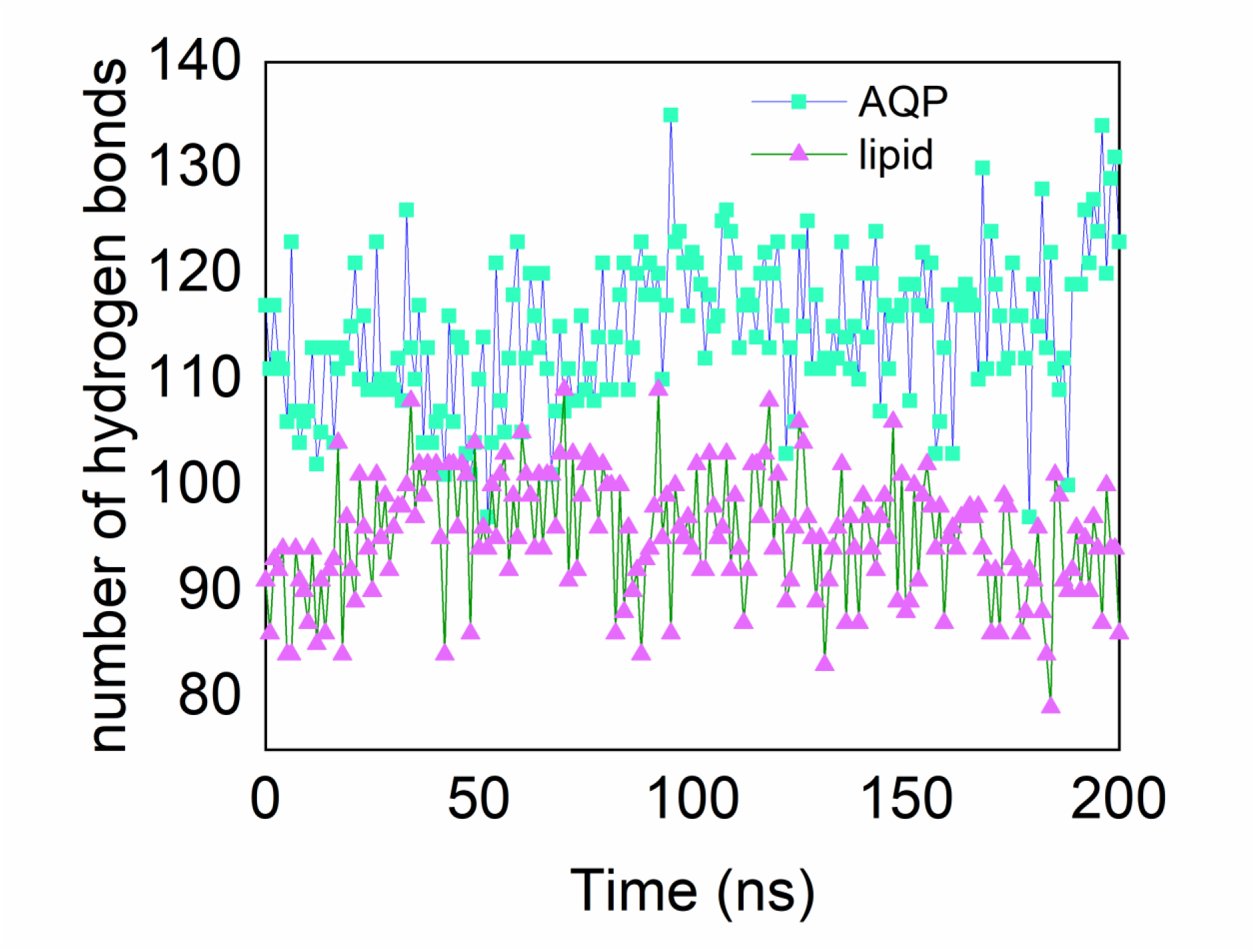
Number of hydrogen bonds formed between peptide and water in lipid and AQP simulation system. Higher number of peptide-water hydrogen bonds were observed when aquaporin is present.

**Figure S3.**
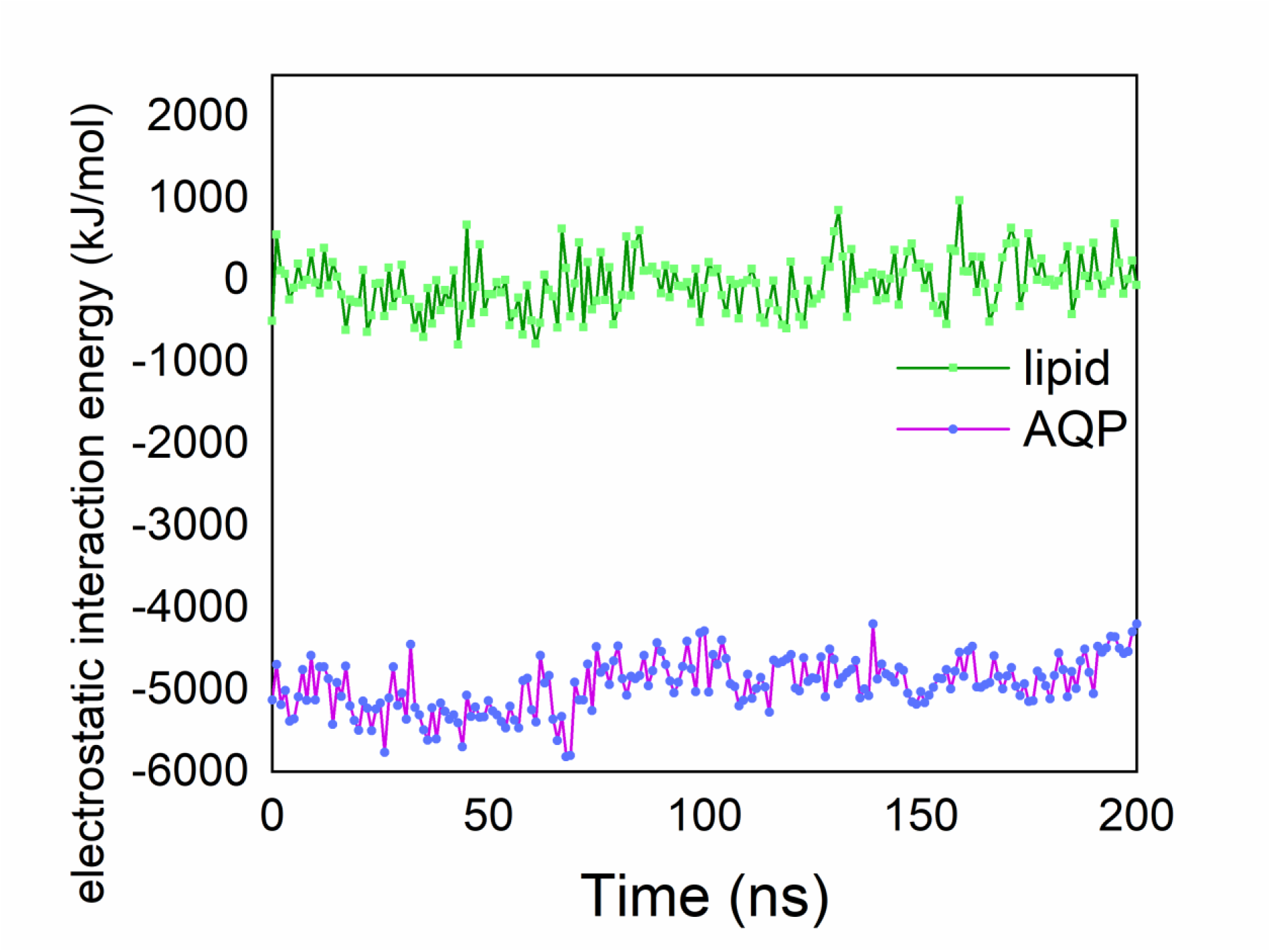
The electrostatic interaction energy between peptide and water. The interaction energy is found to be higher when aquaporin is present.

**Figure S4.**
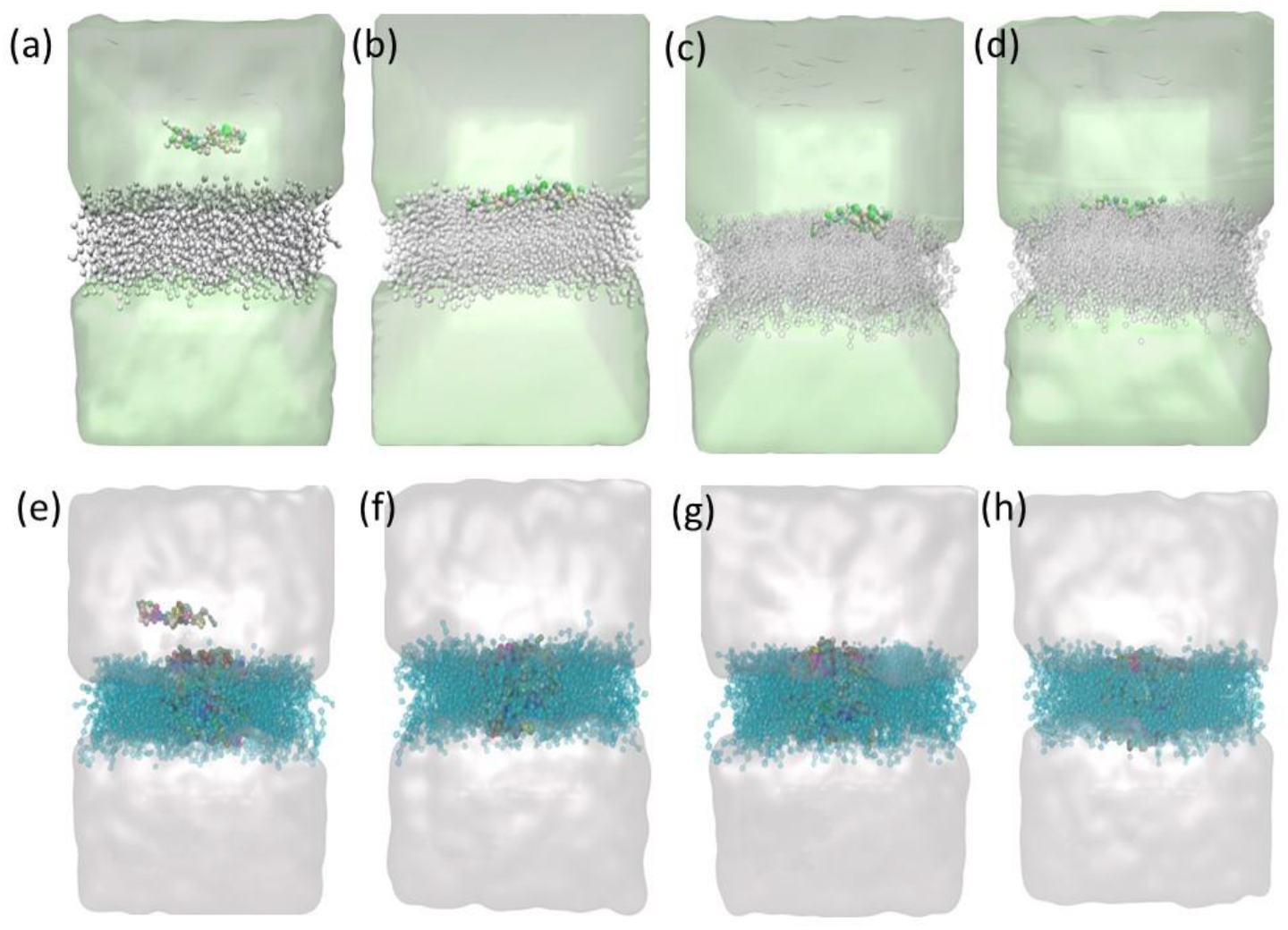
The snapshot from 15 µs CGMD represents the aggregation of amyloid monomers at the surface of the POPC lipid bilayer (top panel) and AQP4 embedded POPC lipid bilayer (bottom panel). The snapshots are at (a)0 µs (b) 5 µs (c) 10 µs (d) 15 µs of the POPC bilayer and (e) 0 µs (f) 5 µs (g) 10 µs (h) 15 µs of the AQP4 surface. Monomers interact with the POPC lipid bilayer and exert detergent-like effects. However, when AQP4 is present, it attracts the pore region of the channel.

**Figure S5.**
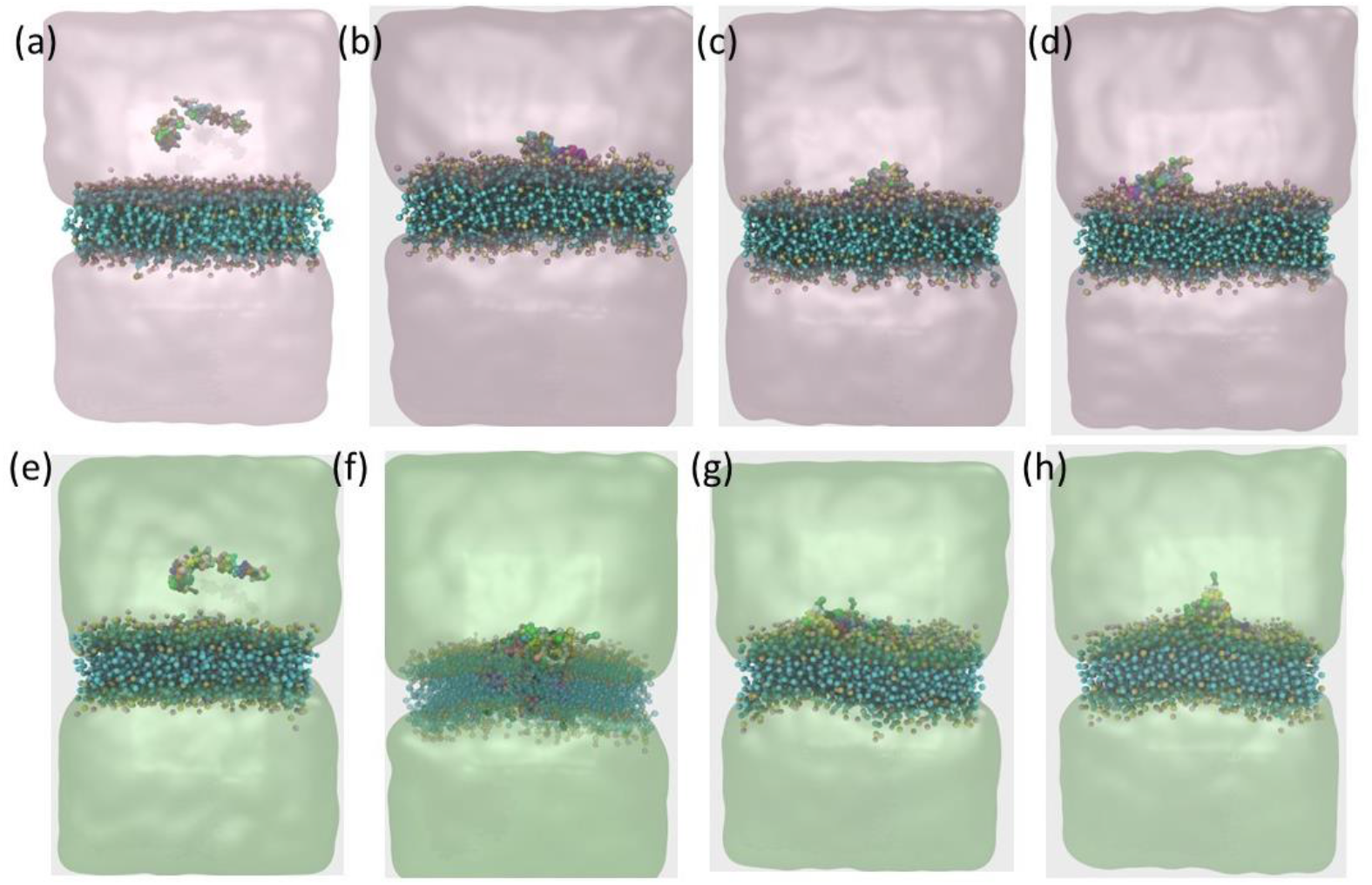
The snapshot from 15 µs CGMD represents the aggregation of amyloid dimers at the surface of the POPC lipid bilayer (top panel) and AQP4 embedded POPC lipid bilayer (bottom panel). The snapshots are at (a)0 µs (b) 5 µs (c) 10 µs (d) 15 µs of the POPC bilayer and (e) 0 µs (f) 5 µs (g) 10 µs (h) 15 µs of the AQP4 surface.

**Figure S6.**
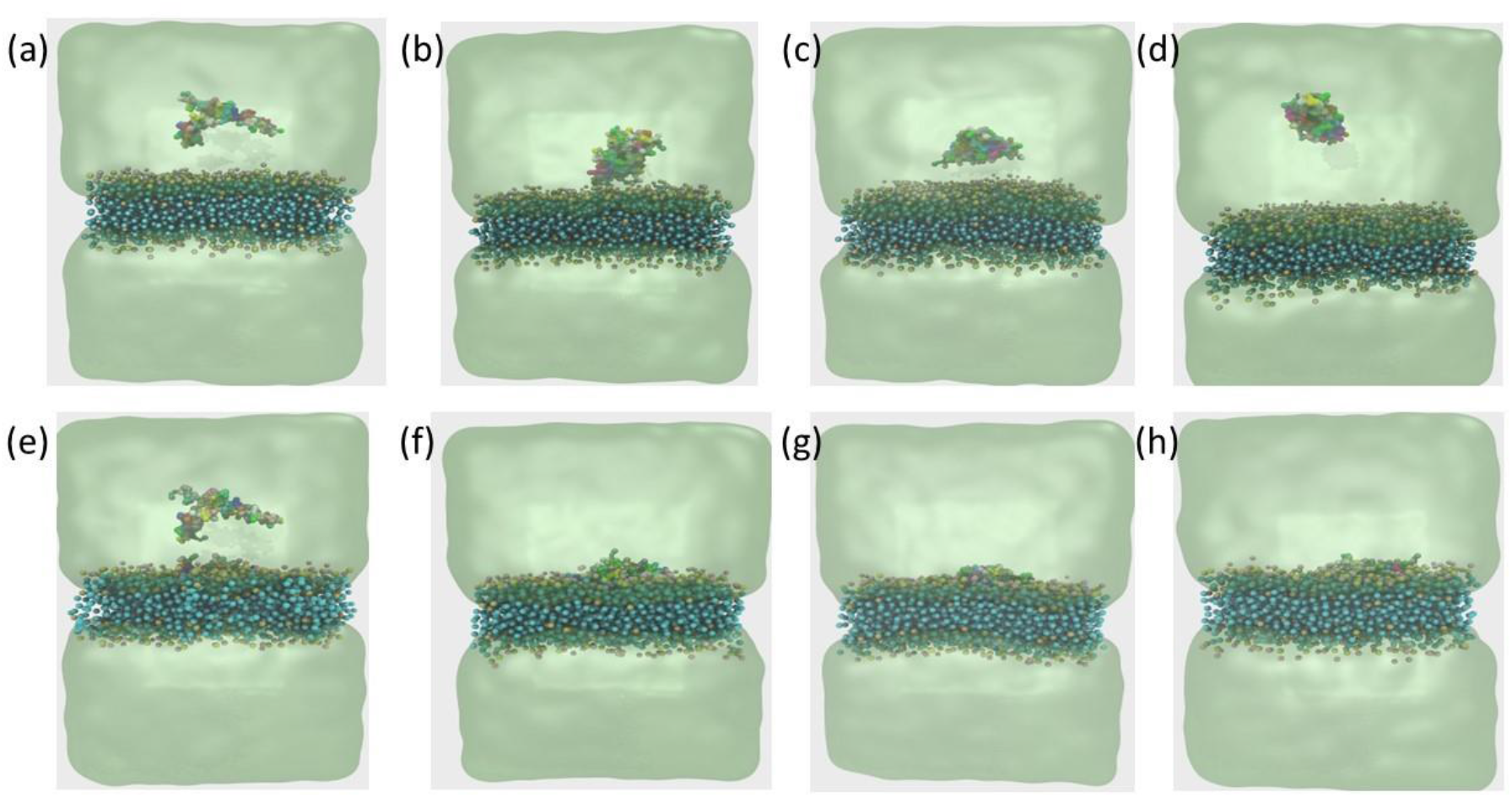
The snapshot from 15 µs CGMD represents the aggregation of amyloid trimers at the surface of the POPC lipid bilayer (top panel) and AQP4 embedded POPC lipid bilayer (bottom panel). The snapshots are at (a)0 µs (b) 5 µs (c) 10 µs (d) 15 µs of the POPC bilayer and (e) 0 µs (f) 5 µs (g) 10 µs (h) 15 µs of the AQP4 surface. The trimers started to interact with water and formed aggregated peptides, which may have been deposited on the surface. However, the presence of AQP4 pulls the peptides towards the pore region and prevents aggregation in water.

**Figure S7.**
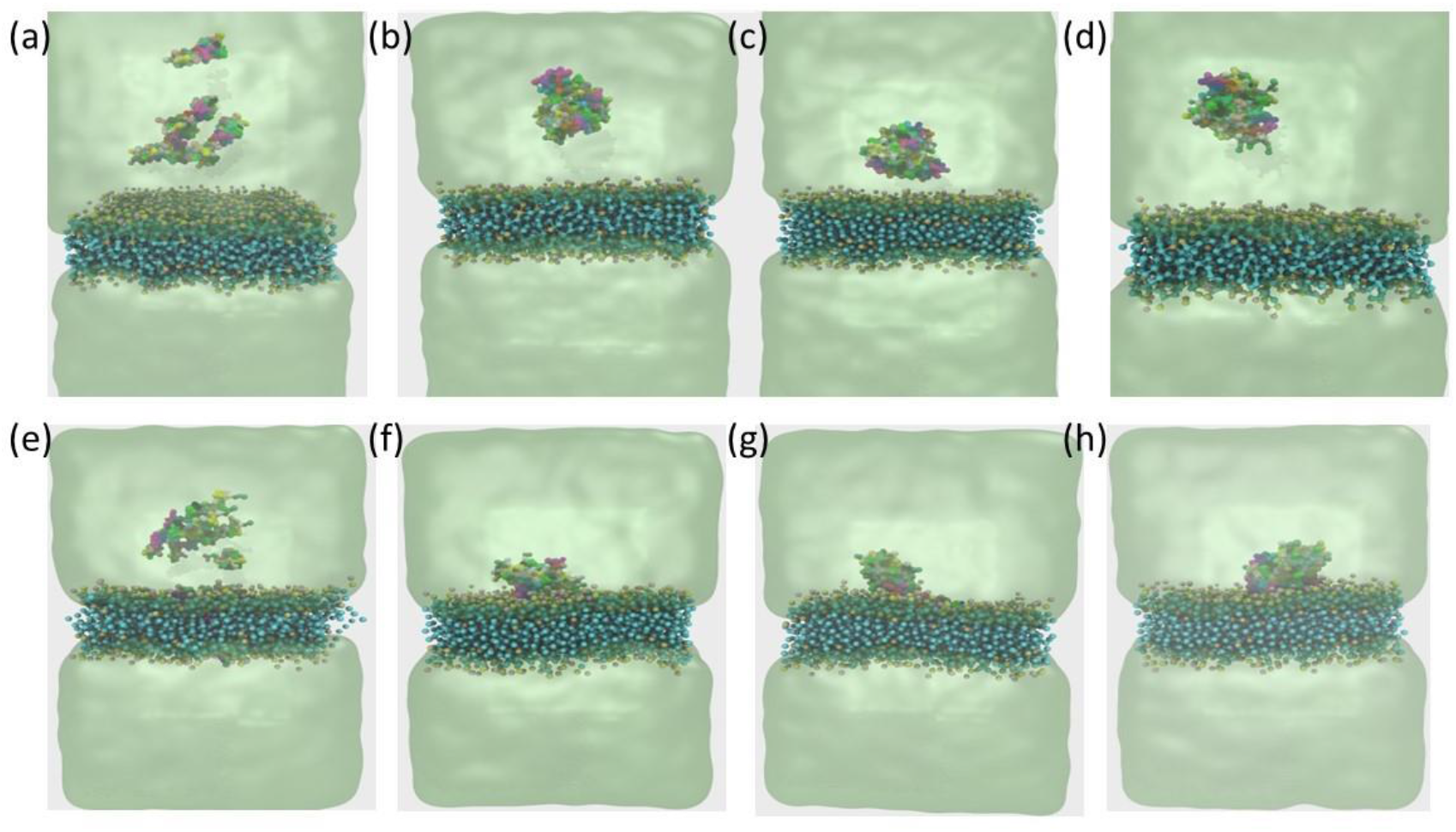
The snapshot from 15 µs CGMD represents the aggregation of amyloid tetramers at the surface of the POPC lipid bilayer (top panel) and AQP4 embedded POPC lipid bilayer (bottom panel). The snapshots are at (a)0 µs (b) 5 µs (c) 10 µs (d) 15 µs of the POPC bilayer and (e) 0 µs (f) 5 µs (g) 10 µs (h) 15 µs of the AQP4 surface.

**Figure S8.**
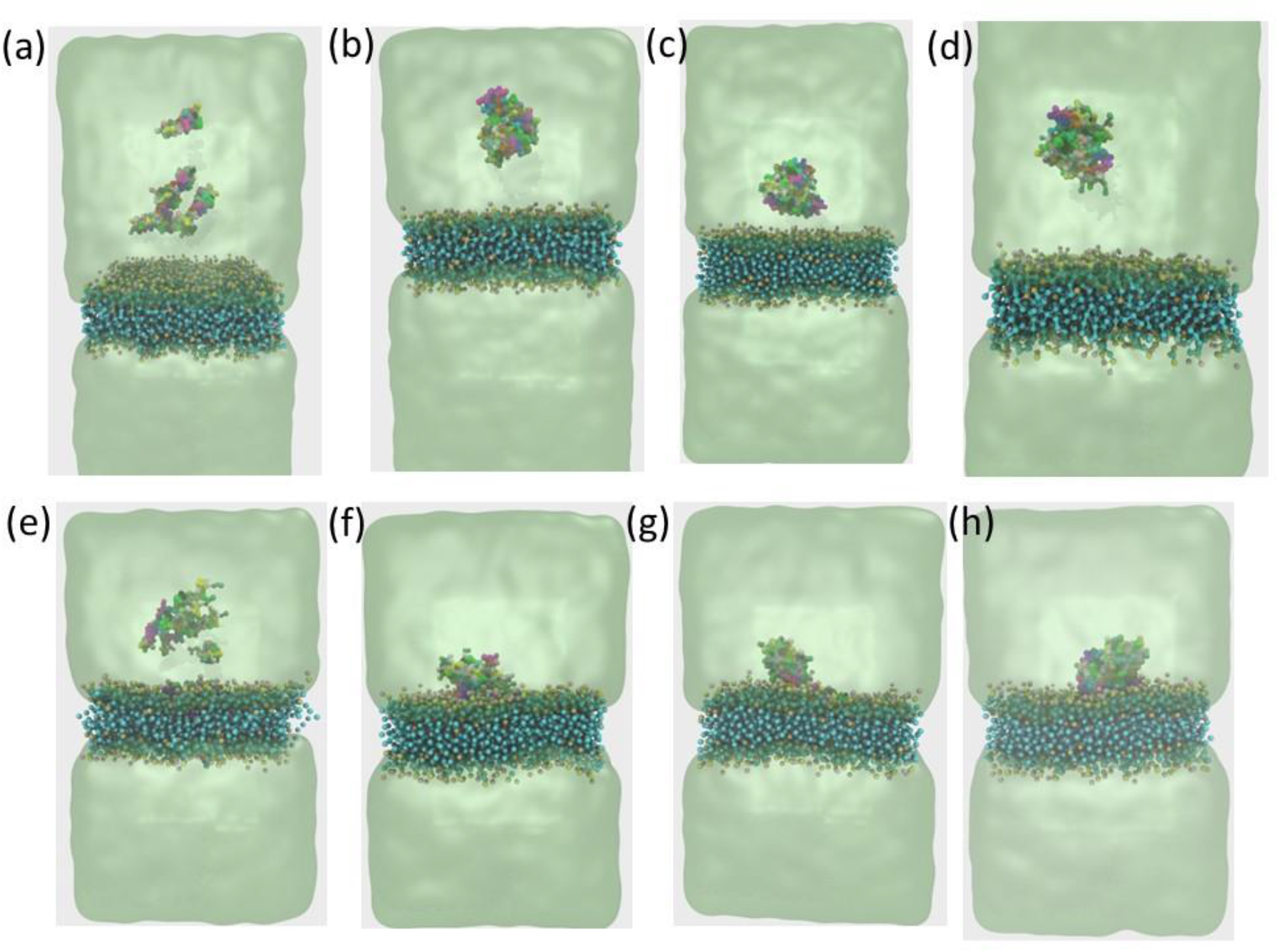
The snapshot from 15 µs CGMD represents the aggregation of amyloid pentamer at the surface of the POPC lipid bilayer (top panel) and AQP4 embedded POPC lipid bilayer (bottom panel). The snapshots are at (a)0 µs (b) 5 µs (c) 10 µs (d) 15 µs of the POPC bilayer and (e) 0 µs (f) 5 µs (g) 10 µs (h) 15 µs of the AQP4 surface.

**Figure S9.**
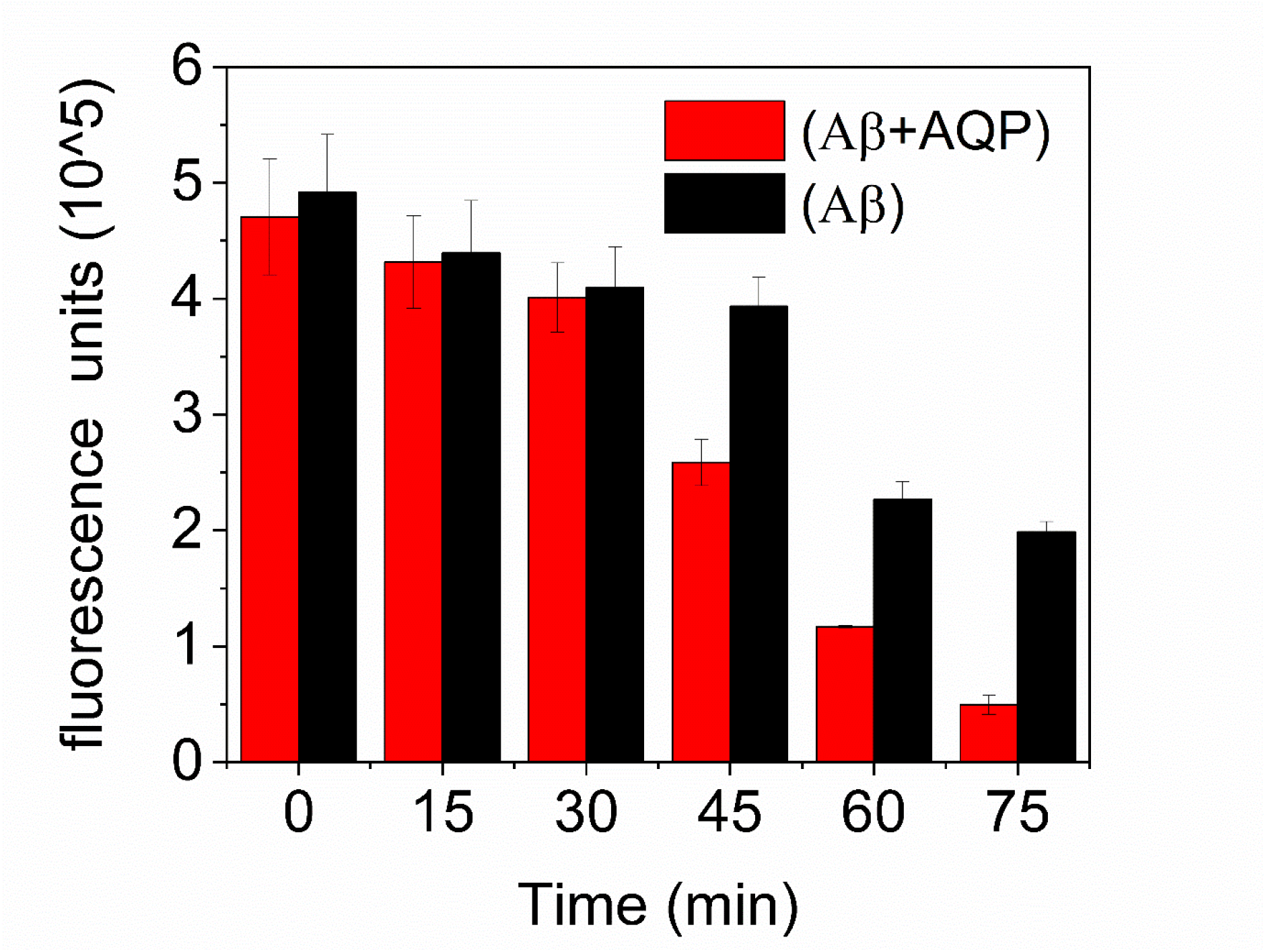
Time-dependent aggregation of amyloid peptides with and without AQP4. Higher fluorescence peaks were observed when AQP4 was absent in the solution.

**Figure S10.**
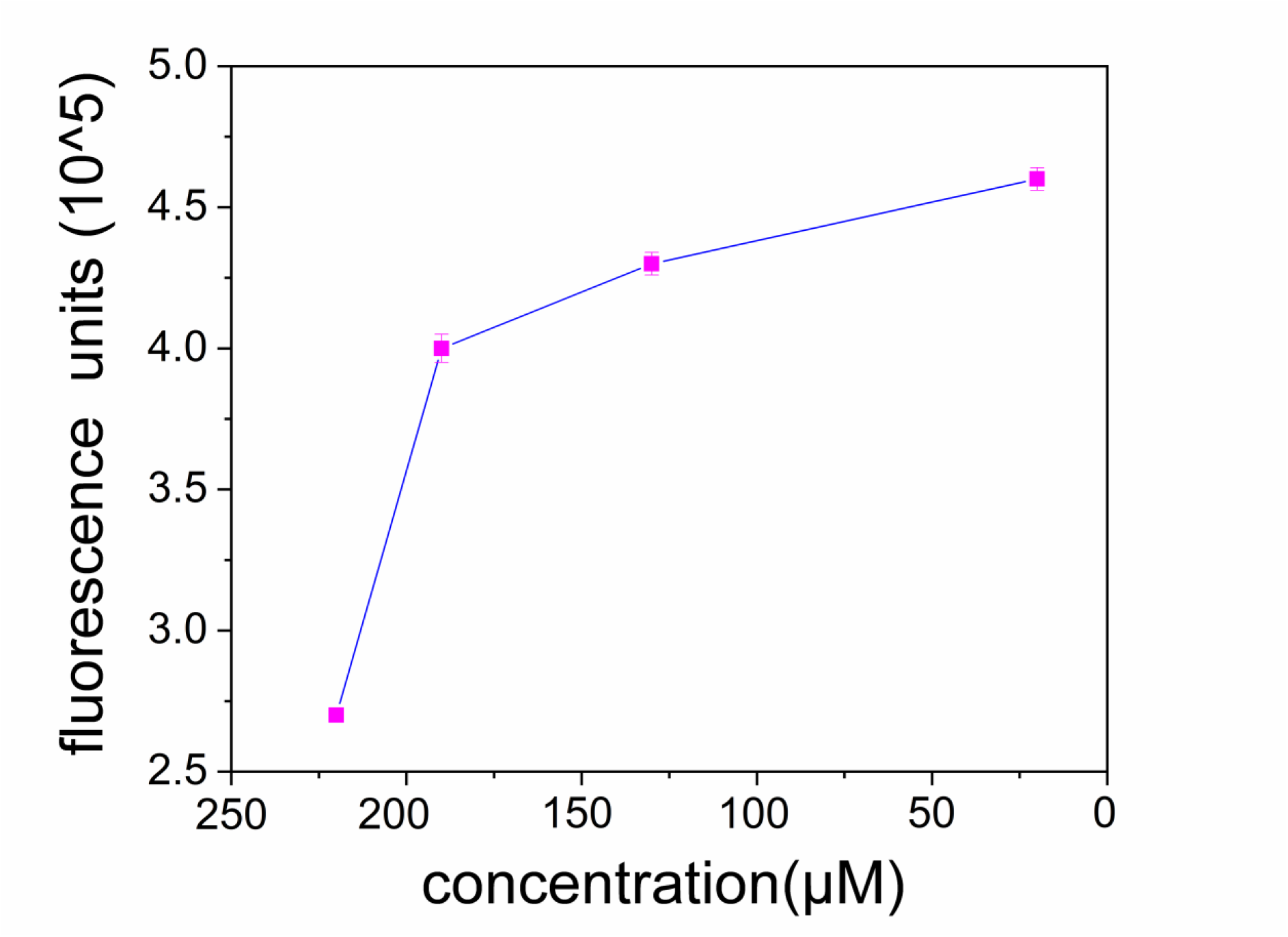
Fluorescence emission of Aβ at varying concentrations of AQP4.

**Figure S11.**
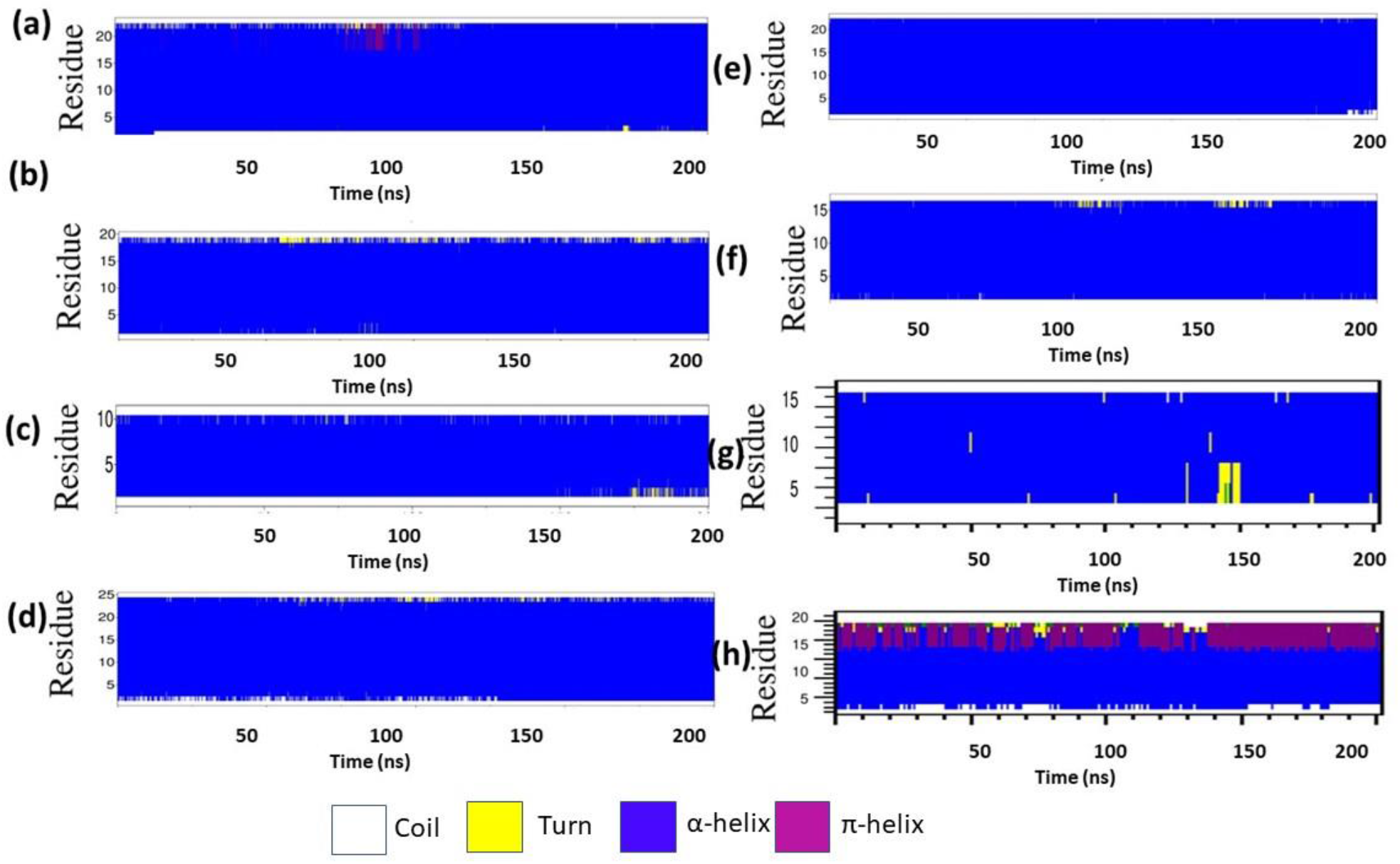
Time-dependent secondary structural changes of helices H1-H8 of the native AQP4 channel. As it can be seen that the helical content is retained during the simulation time.

**Figure S12.**
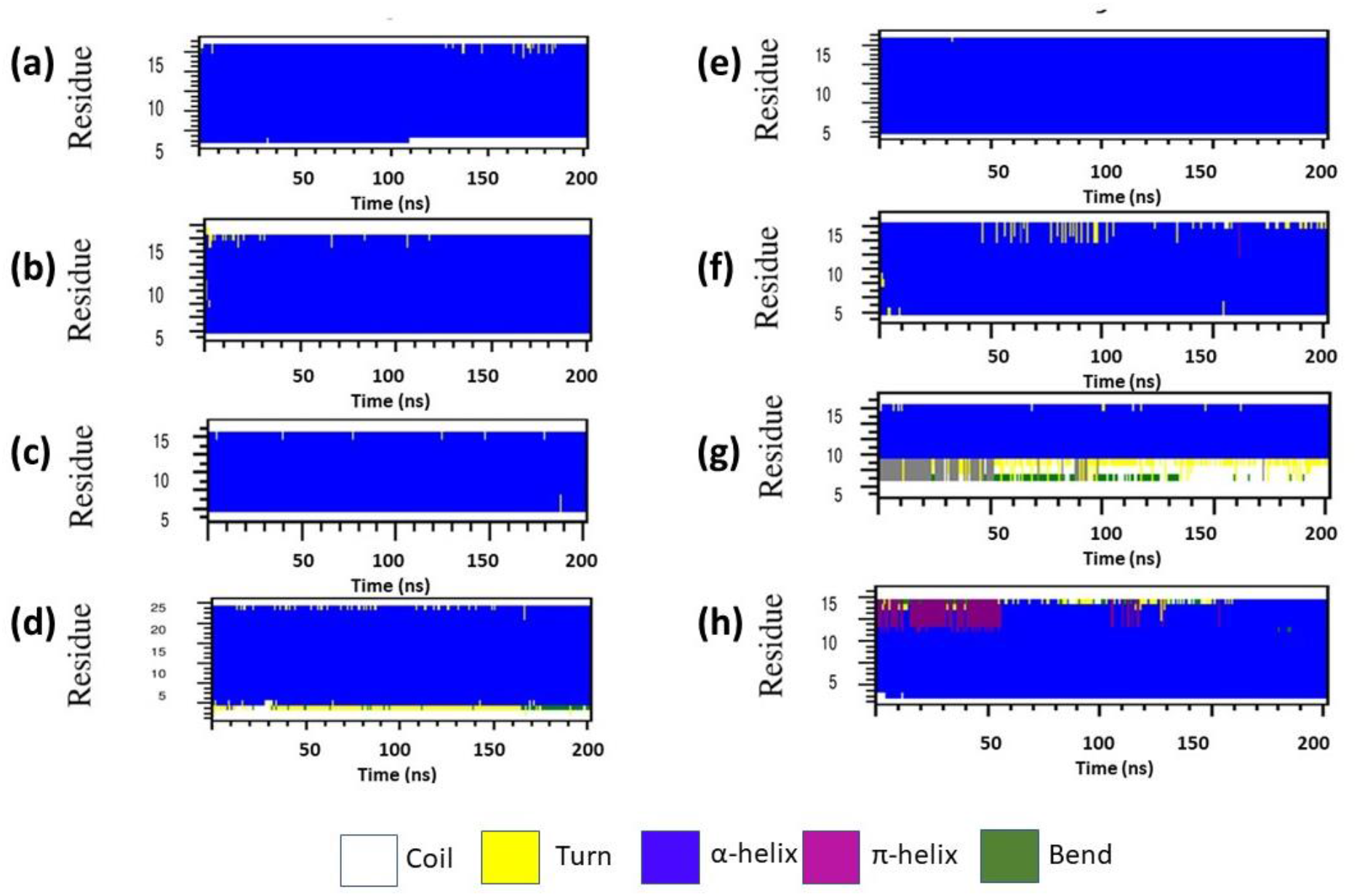
Time-dependent secondary structural changes of helices H1-H8 of AQP4 channel with amyloid monomer. Occasional transitions of helices and formation of turn, coil, and bend regions were observed.

**Figure S13.**
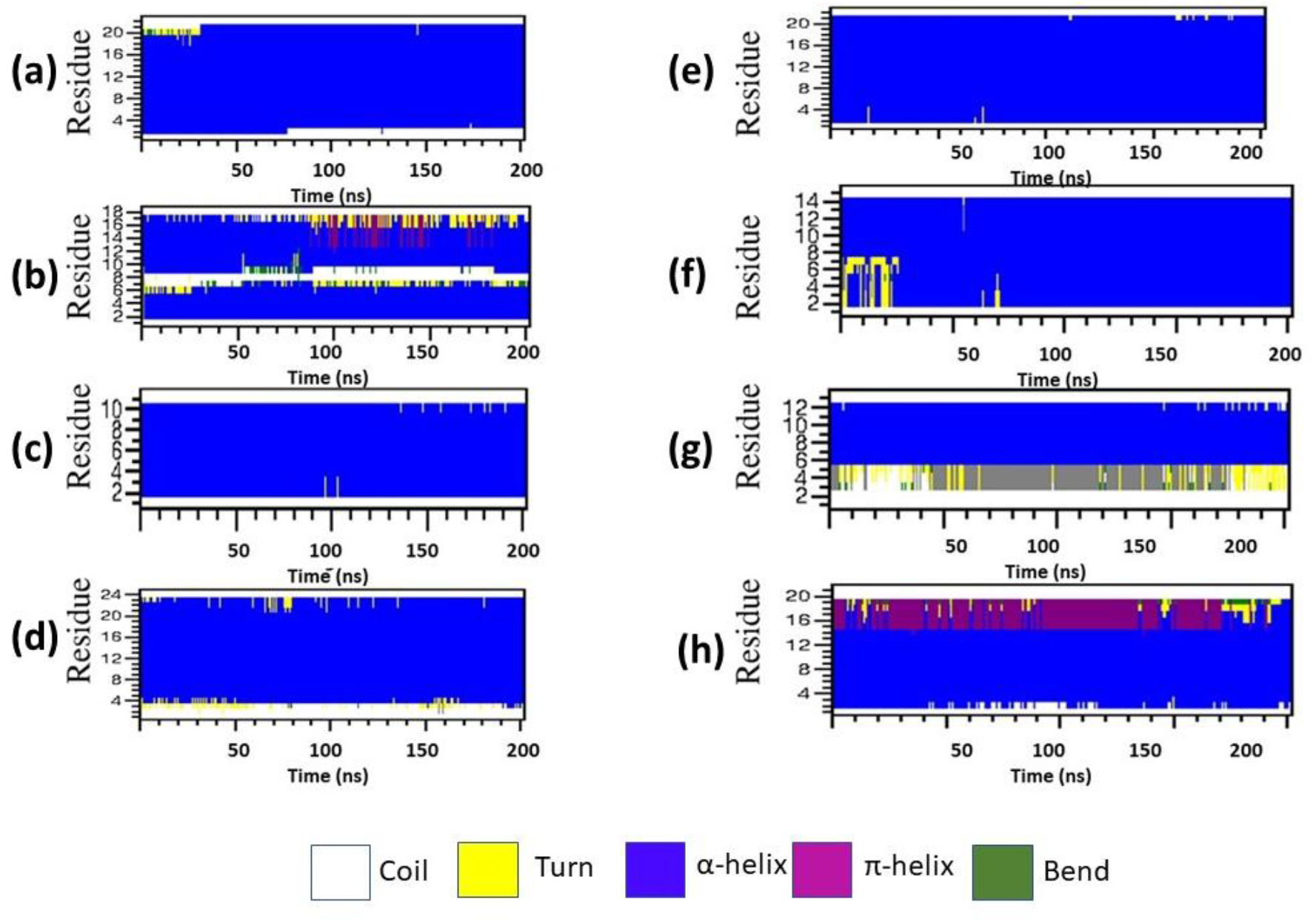
Time-dependent secondary structural changes of helices H1-H8 of AQP4 channel with amyloid dimer. Occasional transitions of helices and formation of turn, coil, and bend regions were observed.

**Figure S14.**
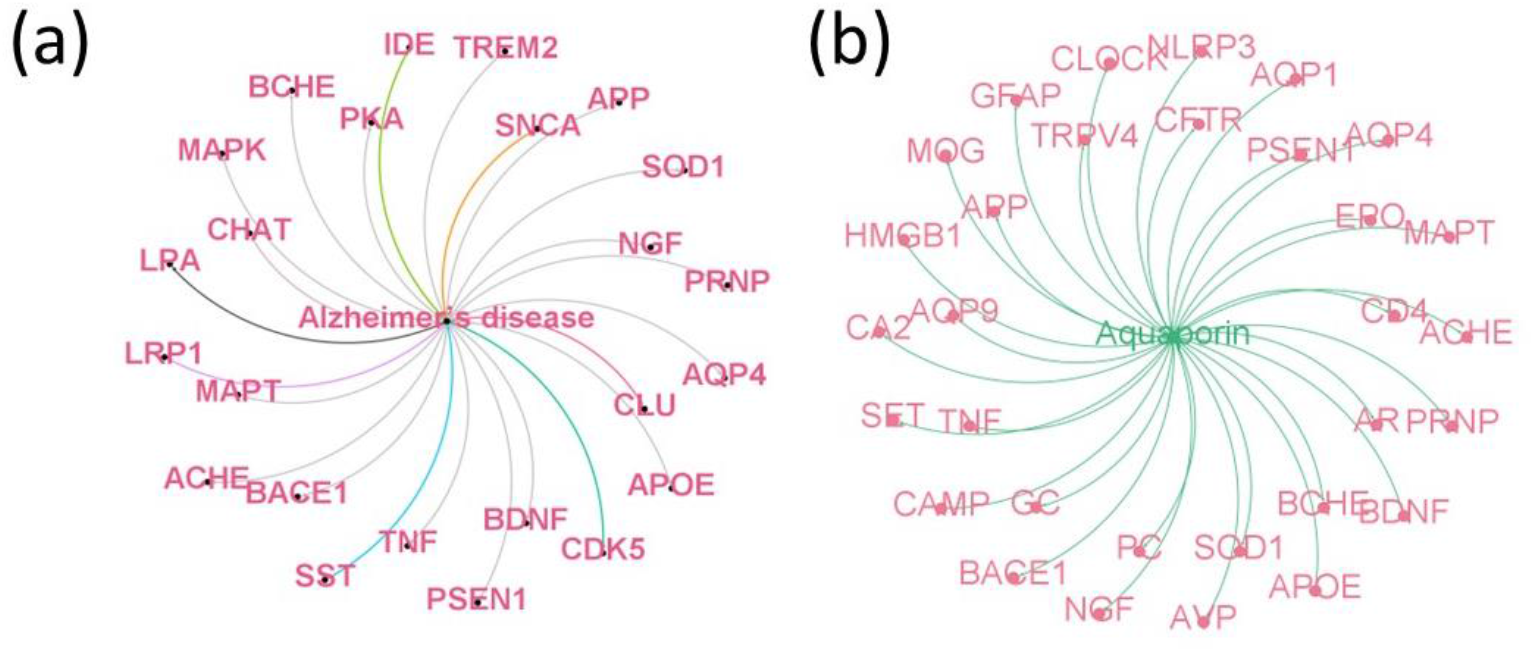
Co-occurring proteins were identified through network analysis using text mining for Alzheimer’s disease and aquaporin-4.

